# Population structure across scales facilitates coexistence and spatial heterogeneity of antibiotic-resistant infections

**DOI:** 10.1101/469171

**Authors:** Madison S. Krieger, Carson E. Denison, Thayer L. Anderson, Martin A. Nowak, Alison L. Hill

**Author notes:** These authors contributed equally to this work (Carson E. Denison, Thayer L. Anderson).

## Abstract

Antibiotic-resistant infections are a growing threat to human health, but basic features of the eco-evolutionary dynamics remain unexplained. Most prominently, there is no clear mechanism for the long-term coexistence of both drug-sensitive and resistant strains at intermediate levels, a ubiquitous pattern seen in surveillance data. Here we show that accounting for structured or spatially-heterogeneous host populations and variability in antibiotic consumption can lead to persistent coexistence over a wide range of treatment coverages, drug efficacies, costs of resistance, and mixing patterns. Moreover, this mechanism can explain other puzzling spatiotemporal features of drug-resistance epidemiology that have received less attention, such as large differences in the prevalence of resistance between geographical regions with similar antibiotic consumption or that neighbor one another. We find that the same amount of antibiotic use can lead to very different levels of resistance depending on how treatment is distributed in a transmission network. We also identify parameter regimes in which population structure alone cannot support coexistence, suggesting the need for other mechanisms to explain the epidemiology of antibiotic resistance. Our analysis identifies key features of host population structure that can be used to assess resistance risk and highlights the need to include spatial or demographic heterogeneity in models to guide resistance management.

## Introduction

Antibiotic resistance is a major threat to our ability to treat bacterial infections. Over the past century, resistance to each new class of drugs has appeared soon after clinical use began. Today, drug-resistant infections are estimated to cost perhaps $20 billion annually^1^. Individual bacteria that are resistant to multiple classes of antibiotics are now common in species such as *Streptococcus pneumoniae, Pseudomonas aeruginosa*, and *Clostridium difficile*^2^, and nearly untreatable strains of *Neisseria gonorrheae*^3^, *Klebsiella pneumoniae*^4^, and *Acinetobacter baumannii*^5^ have recently been identified. These trends have led to speculations about a “post-antibiotic future”, in which routine medical procedures such as surgeries, childbirth, and dental work might become as high-risk as they were in the pre-WWII era due to a lack of effective antibiotic prophylaxis and treatment^6,7^.

Beyond sensational headlines, there is widespread interest among experts in predicting the future morbidity, mortality, and economic impact of drug-resistant infections, so that appropriate investments to counter these trends can be encouraged from government and industry. Mathematical modeling has traditionally played a key role in predicting the dynamics of infectious diseases^8^, and has a long history of application to antibiotic-resistant infections^9^. Despite this, there are several recurrent trends in the spatiotemporal dynamics of drug-resistant bacteria that are difficult to explain. Firstly, for many bug-drug pairs, resistant strains have not completely displaced drug-sensitive ones, but instead coexist stably at intermediate levels, in contrast to predictions of standard infection dynamics and the ecological principle of competitive exclusion (Figure 1i). Secondly, regions which prescribe similar levels of antibiotic can have very different levels of resistance, and these differences persist over time (Fig. 1ii). Finally, even neighboring regions can have large and persistent differences in resistance frequencies, at the scale of neighborhoods all the way up to countries (Fig. 1iii). Since models must, at minimum, be able to explain current trends before being trusted for forecasting, these fundamental disagreements with data have either discouraged efforts to make predictions or led to widespread suspicion of existing predictions (e.g. the Review on Antimicrobial Resistance^10,11^).

**Figure 1.**
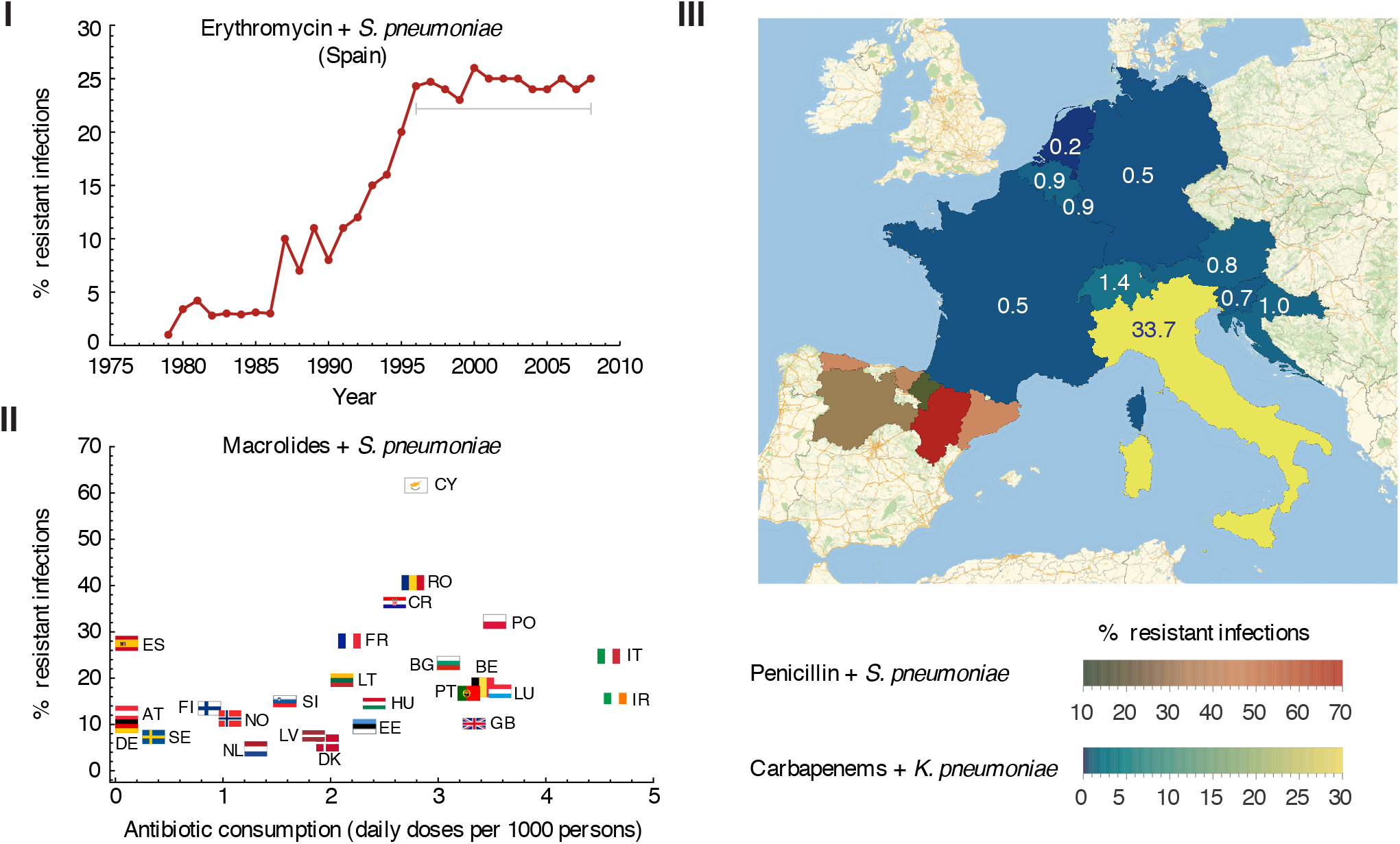
A selection of data illustrating spatiotemporal patterns of antibiotic resistance that common mathematical models have difficulty explaining. **I)** Percentage of *S. pneumoniae* isolates in Spain which are resistant to erythromycin (a macrolide), over time. After an initial increase, drug-sensitive and drug-resistant strains have appeared to coexist stably for ~ 15 years.^12^. **II)** The same amount of antibiotic use can lead to very different levels of resistance in different regions. Percentage of *S. pneumoniae* isolates from 2016 that were resistant to macrolides versus the number of daily doses of macrolides administered per capita, for each country participating in the European Centres for Disease Prevention and Control^13–16^. **III)** Neighboring regions can have vastly different rates of resistance which persist for long time periods. Time-averaged percent of *K. pneumoniae* isolates resistant to carbapenems in Austria, Belgium, Croatia, France, Germany, Italy, Luxembourg, and Slovenia from 2013-2016 and Switzerland from 2015-2016^14,17^ (blue/yellow), along with *S. pneumoniae* isolates resistant to penicillin in six provinces of Spain from 1990-1998 (green/red)^18,19^. The year-to-year deviation from this average is less than 3% (see Fig. S1). Each country is labeled with the resistance level (%). Note that for *S. pneumoniae* and macrolides or penicillin, “% resistant” includes isolates classified as “non-susceptible”.

In this paper we propose a mechanism to explain these perplexing spatiotemporal trends in antibiotic resistance levels that have eluded standard infection models. We consider that host populations may be structured, with heterogeneities in mixing patterns within and between regions, as well as in the distribution of antibiotic use. Spatial differences in treatment rates could arise, for example, from local differences in prescribing guidelines or norms, availability of care, or care-seeking behavior; from the presence of hospitals or other facilities with higher rates of antibiotic usage, or, an increased use of antibiotics following spatially-localized viral epidemics. We find that in contrast to well-mixed populations, structured populations often support long-term coexistence of drug-resistant strains at intermediate levels. We observe that the predicted prevalence of resistance depends not only on the frequency of antibiotic use, but also on how drug use is distributed and details of the transmission network. This mechanism can also reproduce the observation of sharp gradients in resistance levels between neighboring regions, even when infections can spread directly between them. We discuss how these ideas can be used to better understand factors promoting or hindering antibiotic resistance and to construct more realistic models to evaluate antibiotic stewardship policies.

## Observed spatiotemporal trends in antibiotic resistance

### I) Frequencies of resistance over time seem to defy competitive exclusion

For a very large number of bacteria-antibiotic combinations, the prevalence of resistant strains has not reached 100% in the population, but has rather appeared to saturate at an intermediate frequency (Figure 1i). Prominent examples include *Escherichia coli* and aminopenicillins in Europe over the past decade^14,20^, methicillin-resistant *Staphylococcus aureus* in the United States^2^, and penicillin resistance in *S. pneumoniae* in Spain from the 1980’s to 1990’s^18,19^ as well as in other locations/timeframes^21,22^. This long-term coexistence of drug-sensitive and drug-resistant strains is difficult to explain. Resistant strains are clearly selected for in treated individuals, but generally carry a fitness cost such that sensitive strains do better in untreated individuals^23–25^. Mathematical models of infection dynamics under treatment predict that in most conditions the population will move towards an equilibrium where only a single strain persists, with drug-resistant strains reaching fixation when treatment is common enough and costs of resistance are low enough, and drug-sensitive strains dominating otherwise^26^. More generally, long-term coexistence of two strains would seem to defy the ecological principle of competitive exclusion^27–29^, which dictates that when two species compete for a single resource (here, the susceptible population), only one may survive.

Dozens of previous mathematical models have been constructed in an attempt to explain this coexistence. While many studies have focused in particular on *S. pneumoniae*, these and others have attempted to be as general as possible so the conclusions can be extended to other bacterial species. Studies have repeatedly found that well-mixed populations in which individuals can only be infected with one strain (either sensitive or resistant) at a time do not generally support coexistence^26,30–32^. Models that include more details of within-host infection dynamics have suggested some potential mechanisms of coexistence. Drug resistant and sensitive strains can coexist if dual infection of individuals with both is possible, but only for a very narrow range of values for both the treatment coverage level and the cost of resistance^26,30,31^. If *de novo* generation of one strain in an individual infected with the other can occur (presumably via point mutation or horizontal gene transfer from non-pathogenic co-colonizers), then coexistence is again possible^26^, but such a process is likely to lead to low-level mutation-selection balance, not coexistence at near intermediate levels. Recently, Davies et al.^32^ have suggested that co-colonization of hosts with subsequent within-host competition results in a type of frequency-dependent selection that helps maintain coexistence. This occurs because resistant strains have an advantage when they are rare, because they will mostly co-colonize drug-sensitive hosts who, when treated, will be rid of competitors. This mechanism is quite robust to parameter values, but only relevant to commensal bacteria for which co-colonization is common.

Another set of mechanisms shown to promote coexistence is host population structure. If there are two completely isolated sub-populations^26^, or if treated and untreated individuals interact extremely rarely^30^, then sensitive and resistant strains can both persist, but this is unlikely to apply to any realistic scenario. Blanquart *et al*.^33^ recently extended this idea to populations consisting of an arbitrary number of completely interconnected sub-populations, and similarly found that coexistence could occur in some regions of parameter space but only when transmission between the sub-populations was weak. A study simulating transmission in a population in which both contact patterns and the probability of being treated are age-dependent and informed by data for *S. pneumoniae* also reveals more opportunity for coexistence^31^. However, this explanation is still far from general, since coexistence only occurs for costs of resistance <8%, does not display the full range of resistance prevalence levels — and their dependence on antibiotic use — observed in data, and is somewhat reliant on the division of the bacterial population into discrete serotypes which each support different resistance levels.

There are also ecological effects known to promote coexistence of species more generally^34^, but their relevance to bacterial resistance to antibiotics remains unclear. For example, coexistence can occur when species compete more strongly among themselves than with other species^30^. The obvious candidate for such a mechanism in the context of infectious diseases is strain-specific immunity^35^, with the effect that hosts are less susceptible to re-infection by a strain with which they have previously been infected. Strain-specific immunity leads to balancing selection, since low frequency strains have an advantage. However, there is generally not expected to be a connection between the resistance status of a strain and its immunogenicity (e.g. serotype). Lehtinen et al.^36^ have recently shown that such a connection may not be necessary, as linkage between a locus under balancing selection and a polymorphic locus that influences the relative fitness of resistant and sensitive strains (such as duration of carriage for *S. pneumoniae*) can promote coexistence. The relevance of these mechanisms to most antibiotic resistant bacteria remains uncertain.

Overall, despite some progress in explaining the long-term coexistence of drug-sensitive and drug-resistant strains, many proposed mechanisms only reproduce coexistence for small regions of parameter space or species-specific scenarios, and the ultimate set of causes for real-world coexistence is far from fully understood.

### II) Different frequencies of resistance are seen in regions with similar rates of drug prescription

Another puzzling trend in spatiotemporal antibiotic resistance data is the appearance of very different levels of resistance in regions that seem to have the same levels of drug usage. While overall, there is a strong correlation between the amount of antibiotics consumed in a region and the prevalence of drug resistant strains^37,38^, there are many cases that diverge from this trend. For instance, comparing countries participating in the European Centre for Disease Prevention and Control (ECDC), the rate of resistance can vary by as much as 30% even between countries that have equivalent rates of outpatient prescription of penicillin, such as Spain and Poland^13–16^ (Figure 1ii). Other examples can be found at smaller spatial scales, for instance between cities in the American South^39^ and provinces in Spain^19^. While surveillance practices can differ in across regions and nations, the fact that these trends are seen in both ECDC data (where there is significant but not complete standardization) as well as in the smaller-scale studies conducted by single research teams with unified protocols over long time periods, solidifies that they are not artifacts of recording practices.

While this finding has not yet been specifically examined in the context of theoretical models, all of the models mentioned in the previous section^26,30–33,36^ admit only one stable equilibrium for a given set of parameters, including antibiotic consumption level. Therefore, if each region is considered to be an isolated and well-mixed population, extra mechanisms are needed to explain different resistance levels for the same overall antibiotic use. It could be that one or both of the regions in question had not yet reached a steady-state prevalence of resistance by the time of sampling, though when longitudinal surveys of these same resistant strains are available, they generally suggest that levels have approximately equilibrated (e.g. ECDC data^14^). Another possibility is that the cost of resistance varies between regions, for example due to different mechanisms of resistance or different genetic backgrounds (e.g. spa types in *S. aureus*^40^ or serotypes in *S. pneumoniae*^41^), but experimental evidence to support or refute this idea is lacking so far. Finally, the discrete regions considered during resistance surveillance are in reality neither completely isolated nor well-mixed. Movement of individuals and microbes between regions leads to interdependence of resistance dynamics. Within a region, there could be heterogeneous distributions of treatment and infections which impacts the overall resistance level. Consequently, models that attempt to explain this trend must consider the connected nature of regions across spatial scales.

### III) Neighboring regions can show very different frequencies of resistance

Another confounding element of antibiotic resistance epidemiology is the high degree of heterogeneity in resistance levels between neighboring populations at many differing scales. This is slightly different from the preceding point (II): not only do we find two regions with the same levels of antibiotic use but with different levels of resistance, we find that these regions can be bordering each other. Neighboring regions are likely to exchange infected individuals frequently, which would be expected to ameliorate differences in resistance levels over time, just like the predictions of well-mixed models. In contrast, they can actually sustain sharp gradients in the frequency of resistance seemingly indefinitely.

The scales at which this phenomenon is observed range from the size of nations down to neighborhoods of a city (Figure 1iii). For instance, in Europe, the frequency of carbapenems resistance in *K. pneumoniae* has been much much higher in Italy than in neighboring states for many years^14^. Similar trends have led to differences in the frequency of certain resistant strains across the United States^42,43^. At a smaller scale, in the long-term study of *S. pneumoniae* strains in Spain revealed a large difference in frequency between penicillin resistance in Aragon and its neighboring regions^12,18,19,44^. Among municipalities around Copenhagen, Denmark, the prevalence of resistance to trimethoprim and other antibiotics used to treat urinary tract infections in *E. coli* isolates varied by 3-fold^45^. At an even smaller scale, recent (2015) data on the United States revealed that in three neighborhoods in Greater Miami, rates of ciprofloxacin resistance in *Proteus mirabilis* differed by a staggering 62% overall, with rates of 13% (Fort Lauderdale, 172 cases), 41% (Miami, 264 cases), and 75% (Pembroke Pines, 8,979 cases)^39^. The fact that these large differences in resistance levels appear frequently in data at different spatial scales and between neighbors that disburse similar amounts of drug therefore require a unifying explanation.

## Results

### A general model for the evolution of drug resistance in a structured population

To better understand the mechanisms that could be responsible for the spatiotemporal motifs seen in drug resistance data, we developed a simple model for competition between strains of an infection in a structured population. We assume that the total population is divided up into multiple subpopulations (also known as ‘demes’^46,47^) (Figure 2). These demes could represent any subdivision of a human population of interest, such as different countries, regions within a nation, neighborhoods within a city, demographic groups, households, and so forth. Within this population we consider the concurrent spread of drug-sensitive and drug-resistant strains of an infection. Infected individuals spread the disease to uninfected individuals at rate *κ* if they are in the same deme (‘within-deme’ transmission rate) and rate *β* if they are in different but connected demes (‘between-deme’ transmission rate), with *κ* ≥ *β*. We do not allow for any super-infection or co-infection: only susceptible individuals can be infected. Infected individuals recover and become susceptible again at rate *g*.

**Figure 2.**
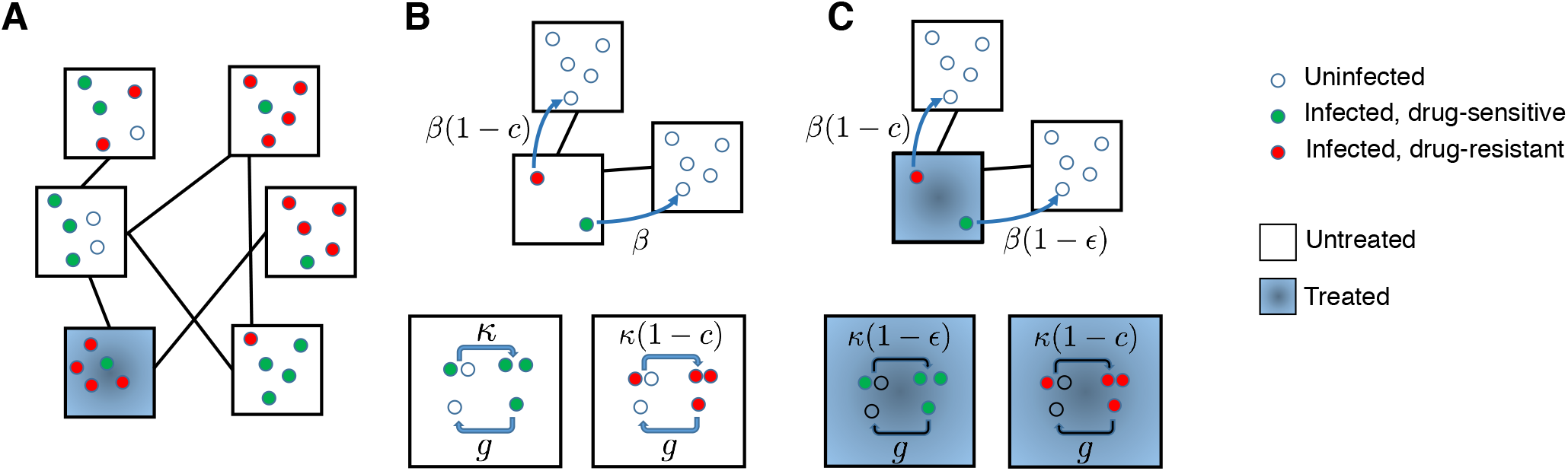
A structured population model for the spread of drug-resistant and drug-sensitive strains of an infection. **A)** An example population, divided into six equal subpopulations (“demes”, black squares) of five individuals (circles). Infection can spread within demes, and also between demes that are connected (black lines). Individuals are categorized based on their infection status (uninfected: open circle, infected with drug-sensitive strain: green circle, infected with drug-resistant strain: red circle). The deme where an individual is located may determine whether or not they will receive drug treatment (blue shading), or more generally, their probability of receiving treatment. **B)** Untreated deme: The wild-type strain (green) is transmitted at rate *κ* within a deme (bottom) and rate *β* between demes (top). Individuals infected with any strain recover with a rate *g*. The resistant strain (red) pays a cost *c* in its transmission rate with or without drug. **C)** Drug-treated deme: Transmission of the resistant strain is unaffected by whether the source individual is receiving treatment, but transmission of the wild-type strain (green) both within (bottom) and between (top) demes is reduced by a factor (1 – *ε*) if the source individual is treated, where *ε* is the drug efficacy.

Each individual has some probability of receiving drug treatment dependent on the deme they belong to. For individuals infected with the drug-sensitive strain, treatment reduces the rate at which they transmit the infection to others (in the same or connected demes) by a factor (1 – *ε*), where *ε* is the efficacy of the drug, 0 < *ε* < 1. We assume that the resistant strain is perfectly resistant to the drug, so that transmission rates of individuals infected with it are unaffected by the presence of treatment. However, the drug-resistant strain incurs a fitness cost *c*, which results in lower transmission rates relative to the wild-type strain with or without treatment: *κ* → *κ*(1 – *c*), *β* → *β* (1 – *c*). Another possibility for the effect of treatment is that it accelerates disease recovery, and we consider this possibility in the **Supporting Information**.

The “infected” status in our model could represent a symptomatic disease state caused by a pathogenic microbe, or an asymptomatic “colonized” state due to a commensal organism. Like others^26,30–33,36^, we do not explicitly model virulence or mortality. Our model is also agnostic as to whether treatment is received only in response to infection (e.g. if a symptomatic infection triggers administration of antibiotics) or independently of infection status (e.g a commensal colonizer is subject to bystander selection from antibiotics administered for another condition^48,49^), since we assume treatment does not impact susceptibility to catching an infection.

Our model is an extension of the classic Susceptible-Infected Susceptible (SIS) model for disease transmission^50^ to a case with two disease strains (sometimes called an *SI*_1_*I*_2_*S* model^51^). We assume that each deme is large enough that the dynamics can be considered deterministically. This model is structurally neutral^52^, meaning that if two identical strains are simulated (e.g. if *ε* = 0 and *c* = 0), they will continue to persist *ad infinitum* at the same level at which they were initiated.

### Absence of coexistence in an unstructured population

We first consider the dynamics of competition between drug-resistant and drug-sensitive strains in a single large, well-mixed deme (an unstructured population) (Figure 3A). A fraction *f* of all individuals infected by either strain will receive drug treatment for the duration of infection. The dynamics are described by a set of four differential equations, tracking the proportion of total individuals in each of four infected states (drug-sensitive untreated, drug-sensitive treated, drug-resistant untreated, drug-resistant treated), while the remaining individuals are uninfected (see **Methods**).

**Figure 3.**
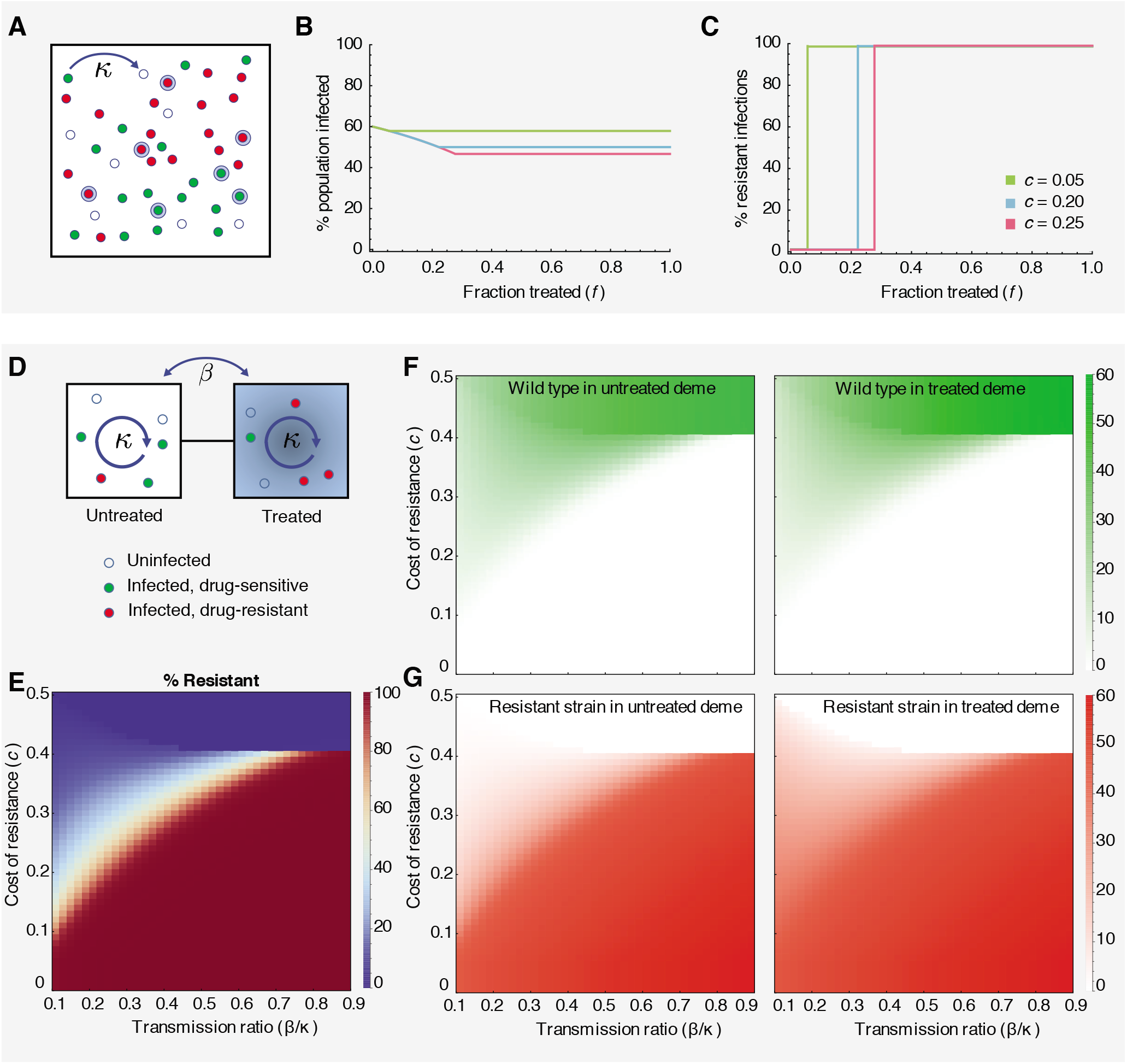
Competition between drug-sensitive and drug-resistant strains in a single well-mixed population (A-C) and two interconnected subpopulations (D-G). **A)** A population of individuals in a single well-mixed deme, in which a fraction *f* receive drug treatment (blue haloes). Individuals may be uninfected (hollow blue circles), or infected with either the wild-type (green circles) or drug-resistant (red circles) strain. **B)** The total prevalence of infection (wild-type + drug-resistant) as a function of the fraction of treated individuals (*f*) for different costs of resistance (*c*). **C)** The % of infections that are drug-resistant as a function of the fraction of treated individuals (*f*) for different parameters. Infection switches between 0% and 100% resistant when *f* = *c*/*ε*. Coexistence never occurs. **D)** Schematic of a two-deme population (left-untreated, right-treated) and the two strains (green-wild type, red-resistant) considered in the model. **E-G)** Each panel shows the infection level (shading) as a function of the relative connectivity between demes (*β* / *κ*) and the cost of resistance (*c*). **E)** The % of all infections that are drug-resistant strain across the entire population. **F)** The % of individuals in each deme who are infected with the wild type strain. **G)** The % of individuals in each deme who are infected with the resistant strain. For all results, the transmission rate is *κ* = 0.25/day, the recovery rate is *g* = 0.1/day, and the treatment efficacy is *ε* = 0.9.

The basic reproductive ratio (*R*_0_), defined as the average number of secondary infections caused by a single infected individual in an otherwise uninfected population, can be defined for each disease strain in this model and completely determines the equilibrium behavior (details in **Supporting Information**). If both infections have *R*_0_ < 1, then neither strain of infection will persist and the equilibrium consists only of uninfected individuals. Otherwise, only the strain with the larger *R*_0_ will persist (at a prevalence 1 – 1/*R*_0_, Figure 3B) while the other will go extinct. Based on the formulas for *R*_0_, the resistant strain persists if and only if the coverage and efficacy of treatment are enough to offset the cost of resistance, *ε f* > *c* (Figure 3C). Therefore, when the population is well-mixed, the heterogeneity introduced by partial treatment coverage (0 < *f* < 1) is insufficient to allow coexistence of wild-type and drug-resistant infections, and competitive exclusion always occurs, in agreement with other studies^26,30–32^.

### Limited coexistence in two connected subpopulations with heterogeneous treatment distribution

We next consider the case of two separate but connected subpopulations of equal size (Figure 3D). As an extreme example, we assume that individuals in one of the demes always receive treatment, while individuals in the other deme are never treated. The system is described by a set of equations tracking the proportion of individuals of each infection status (uninfected, infected with drug sensitive, infected with drug resistant) in each deme (treated vs untreated). The basic reproductive ratio can be derived for each strain (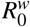 and 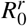) (see **Methods**).

This population supports qualitatively different infection dynamics than the single-deme case (Figure 3E-G). While most parameter regimes still lead to persistence of only the drug-sensitive or the drug-resistant strain (even when *R*_0_ > 1 for both), there is now a stable equilibria where there is a mixture of both strains coexisting (Figure 3D). This region of coexistence occurs when the between-deme infection rate is relatively low compared to the within-deme rate (*β*/*κ* < 0.7 when *ε* = 0.9) and for intermediate values of the cost of resistance *c*. The mathematical derivations of these boundary regions and their physical intuition is included in the **Supplementary Information**. Interestingly, in this mixed equilibrium, both demes contain a mixture of sensitive and resistant strains, a stricter type of strain coexistence than a situation in which each strain dominates a different deme.

This simple example, consisting of only two preferentially-mixing subgroups of equal size, demonstrates that co-clustering of transmission and propensity for antibiotic use can lead to stable coexistence of sensitive and resistant strains at not only the population level but also the *sub*population level, suggesting a resolution for the ubiquity of coexistence in empirical data (observation **(I)**). This host population structure produces a situation in which even though a strain is disfavored in a global sense by having a smaller *R*_0_ value than its competitors, it can be locally favored and thus avoid extinction. However, in this simple example, the region in parameter space where coexistence is seen is relatively small, even for this extreme example of treatment clustering, motivating the question of whether more complex structures will expand the stability of coexistence. Moreover, this two-deme example cannot explain the observation that regions receiving the same amount of treatment or regions neighboring one another often support very different levels of resistance (observations **II** and **III**). Replicating real-world data requires finding situations in which neighbors with *the same* level of drug treatment still have very different levels of resistance. In the subsequent sections, we examine whether more complex population structures can recreate these observed patterns.

### Robust coexistence in more complex population structures

Our results for simple one- and two-deme scenarios suggest that host population structure can promote coexistence of drug-sensitive and resistant strains of an infection. To investigate this relationship further, we extended our model to include an arbitrary number of demes and connectivity patterns (Figure 2A). For simplicity, we first assumed that each deme in the population was the same size and was connected randomly to a fixed number of other “neighbor” demes, therefore creating a collection of random regular graphs^53^ (Figure 4A). Although in reality much more complex human population structures are possible, these simplified networks provide a good base to examine the impact of specific properties on infection dynamics. Each deme was independently assigned to be “treated” (meaning individuals in that deme receive treatment, while individuals in “untreated” demes do not). We relax these assumptions in later sections.

**Figure 4.**
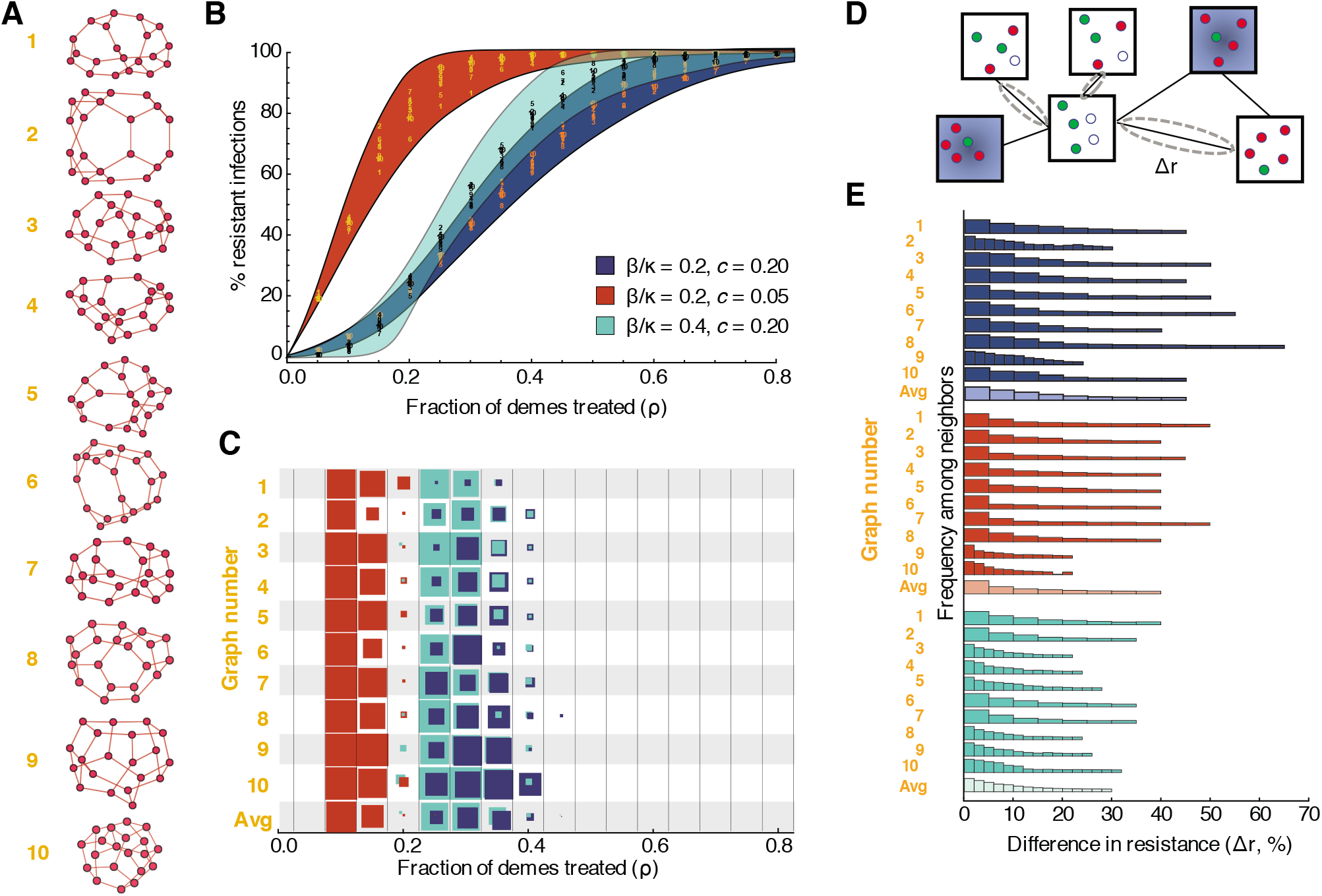
Dynamics of drug-resistant infections in populations consisting of networks of inter-connected demes. **A)** Randomly generated population structures on which infection was simulated. Each node represents a deme (a well-mixed sub-population of individuals), and each edge indicates that infection can spread in either direction between those two demes. Ten example populations were selected out of 1000 total simulated, each with twenty demes randomly connected to three neighbors each, to represent a broad range of outcomes. **B)** Fraction of infections that are resistant in the entire population (*y*-axis) versus fraction of demes treated, *ρ* (*x*-axis). Each color represents a different parameter set (blue background — baseline, red background — lower cost of resistance, teal background — more between-deme connectivity). Numbers show data points for the ten example populations. The colored envelope is created by shading between sigmoidal curves that encompass all the data. **C)** For each population structure shown (*y*-axis) and each treatment level (*x*-axis), the proportion of simulations that resulted in *robust* coexistence between drug-sensitive and drug-resistant strains is shown (by the colored area of the box). Robust coexistence was defined as at least 80% of demes supporting both strains at frequencies above 10%. **D)** Differences in resistance levels (% of all infections that are with the drug resistant-strain) are measured between all pairs of directly-connected untreated demes. **E)** Histograms showing the distribution of pairwise differences in resistance for a given population structure. Lighter shaded histograms combine results from all population graphs. All simulations used kinetic parameters *κ* = 0.25/day, *g* = 0.1/day, and *ε* = 0.9, and pooled results from 100 simulations with different random allocation of treatment across demes. Pairwise differences were calculated with 30% treatment.

Using these “meta-populations”, we found that coexistence is possible for a broad range of costs of resistance, treatment levels, and population structures. As observed in data, the overall prevalence of resistance in a population is roughly correlated with the treatment coverage, increasing gradually as treatment levels increase (Figure 4B). Moreover, when resistance levels were intermediate, coexistence also persisted at the level of individual demes. We defined ‘robust’ coexistence to mean that at least 80% of demes had at least 10% of infections caused by each strain. For the baseline parameter values we used a 20% cost of resistance, treatment levels between 20 and 40% supported robust coexistence in at least some population structures and treatment allocation schemes. For a lower cost of resistance (*c* = 0.05), robust coexistence occurred at a lower range of treatment levels (Figure 4C). When the amount of infection spread (*β*) arising from contact between “neighboring” demes increases relative to the within-deme spread (*κ*), resistance increases more sharply with treatment level, but regions of robust coexistence remain (teal color, and more parameter values in Figures S14 and S15).

These results show that even relatively simple host population structures can reconcile mathematical models with empirical observations of long-term coexistence between drug-sensitive and drug-resistant strains. These results also provide a connection between our model and the data for another spatio-temporal observation, **(II)**: the same overall rate of drug consumption can lead to different prevalence of drug resistance in different populations (even if treatment efficacy and cost of resistance are the same). For example, in Figure 4B, a particular treatment level (e.g. 20% for the red parameter set) can be associated with very different frequencies of resistance in different population structures (up to 30% difference between graph 6 and 9, similar to the data in Fig. 1). A more detailed study of the role of higher-order network properties is presented in a later section.

### Large differences in resistance levels are possible even between connected regions

With these complex multi-deme populations, we examined whether neighboring regions with equal levels of antibiotic consumption could sustain vastly different amounts of resistance (observation **(III**)). For each population structure, random treatment allocation, and parameter set, we chose all nearest-neighbor pairs which were both untreated, and generated a distribution of the pairwise differences in resistance levels between these neighbors (Figure 4D,E). We found that large differences in resistance levels between neighbors were common, with 34% of pairs receiving no treatment differing by more than 10% resistance prevalence, for our baseline parameter values when averaging across all population structures (Figure 4E). In the same simulations, 2% of pairs overall – and up to 15% in some structures – differed by greater than 30% resistance frequency, a value near the upper limit of observations from data shown in Figure 1. Differences of up to 60% between neighbors were observed in some individual simulations.

While populations with higher levels of between-deme mixing (higher *β* / *κ*, teal color) often supported persistence of both drug-sensitive and drug-resistant strains at equilibrium (Figure 4B), they supported smaller differences in resistance levels between demes receiving the same treatment (Fig. 4E). In this parameter regime, sensitive and resistant strains are more segregated between untreated and treated demes, respectively, explaining why simulations on these structures rarely produced “robust” coexistence as we defined it (over 80% of demes supporting at least 10% of each strain) (Figure 4C). We also found that populations with weak connections between all demes were less likely to have large pairwise differences in prevalence of resistance (Fig. S11). Overall, these findings corroborate the intuition that adding edges between demes or raising the between-deme infection rate of connected demes brings the system closer to well-mixed population and hinders coexistence.

### Properties contributing to higher resistance

In order to understand how specific properties of host population structure contribute to the frequency of resistance and the likelihood of coexistence, we simulated infection dynamics on a large set of transmission networks and then statistically analyzed the results. To first generate a large ensemble of population structures which varied in many graph-theoretic properties, we used the Watts-Strogatz^54^ algorithm to create 1000 unique networks of 50 demes each, which had different average degree, variance in degree, clustering, centrality, efficiency, and mean path length (see **Methods** for definitions of these properties and Figure S3 for their values). For each graph, we generated 50 different allocations of treatment, keeping the overall population treatment level the same. Then, we collected the results and employed LASSO (least absolute shrinkage and selection operator) regression to identify the most important properties contributing to the frequency of resistant infections (Table 1). The regression was conducted both at the level of individual demes (i.e. to identify local properties of demes that influenced the level of resistance within a deme) and at the level of the whole population (i.e. to identify properties of the whole population that influenced the overall level of resistance). The simulations and the regression were repeated for a “low resistance” case, in which overall 24% of individuals in the population were treated and as a result less than half of infections were resistant at equilibrium, and a “high resistance” case, where 40% were treated and over half of infections were resistant (Figure S4). Properties were classified based on whether they were always associated with an increase or decrease in resistance, or whether they pushed resistance towards intermediate levels (i.e. facilitated coexistence) or towards extreme levels (i.e. hindered coexistence).

**Table 1.**
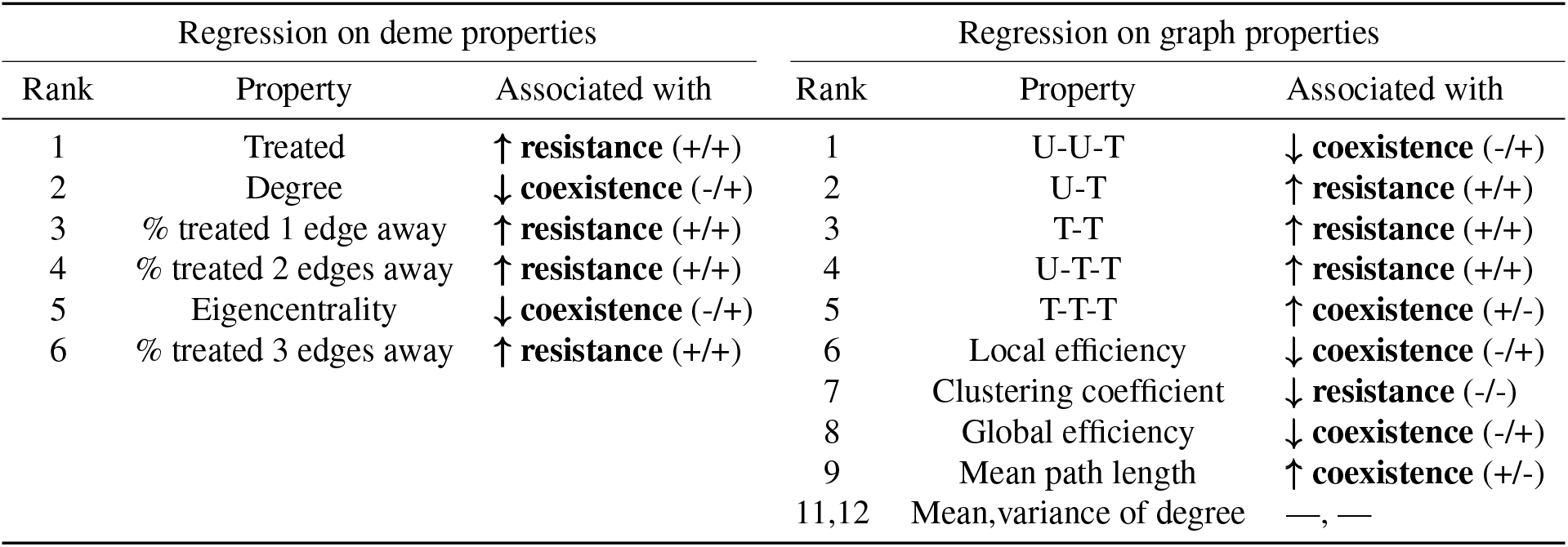
Properties of demes and graphs that accurately and sparsely predict the frequency of drug-resistant infections according to LASSO regression. After running thousands of infection simulations with varying transmission networks and treatment allocations, we used LASSO regression to to determine the association of each property with the prevalence of resistant infections. Regression was conducted separately at the deme (left) and graph (right) level and properties were ranked based on their predictive strength (see **Methods**). For each regression, the sign of the direction of the relationship between the property and the frequency of resistance was determined (+ or -), and used to summarize the associations: “↑ resistance”: (+/+) the property is always associated with increased resistance, “↓ resistance”: (-/-) the property is always associated with decreased resistance, “↑ coexistence”: (+/-) the property is associated with increased resistance when resistance is rare but decreased resistance when resistance is common, “↓ coexistence”: (-/+) the property is associated with decreased resistance when resistance is rare but increased resistance when resistance is common, “—”: the property is not used in the “best” (error-minimizing) model. The shorthand with “U” and “T” indicates the frequency of pairs and triplets (including triangles) of **<u>u</u>**ntreated and **<u>t</u>**reated demes; thus, U-T-T indicates the frequency of three connected demes of which two are drug-treated, normalized by the number of all three-deme combinations in the graph.

We first examined predictors of the resistance level within a deme (Table 1, left side). The most important contributor to resistance was whether or not the deme itself was treated, and the distribution of treatment among the deme’s first, second, and third nearest neighbors were also important – albeit less so – predictors. The next strongest predictor was the total number of first degree neighbors of a deme (the “degree”). Higher degree is associated with lower resistance when resistance is rare, but higher resistance when it is common, and hence pushes resistance to more extreme levels, acting against coexistence of both strains within a deme (−/+ in Table 1). The same is true for the eigencentrality of the deme, a measure of its overall level of connectivity within the network. These results imply that highly-interconnected, hub-like demes are more likely to harbor the more common strain, while demes that are on the periphery of a network are more likely to support coexistence of the strain that is rarer in the population overall. Hence, when resistance is rare, it may be preferentially found in less connected demes, whereas when it is more common, it may cluster in more central demes.

We next evaluated predictors of resistance at the level of the entire population (Table 1, right side). Surprisingly, the simplest statistics of a graph – including the mean and variance in the number of neighbors of each deme – were the least important contributors to resistance. Rather, the best predictors involved a mix of treatment-clustering properties, like the proportion of sets of three connected demes where two of the demes were treated (“U-T-T”), and node-clustering properties, such as measures of “efficiency”. Treatment promotes consistently higher resistance levels when it is distributed more evenly throughout the population (e.g. treated and untreated demes are interspersed in the network, “U-T-T”, “U-T”), but more extreme clustering of communities receiving the most treatment (“T-T-T”) instead facilitates coexistence. This suggests that the effect of increasing antibiotic consumption on resistance levels will depend strongly on which communities experience the increases. The clustering coefficient and local efficiency are both measures of how interconnected the neighbors of a given deme are, on average. Global efficiency and mean path length are both measures of the ease of moving between any two random demes in the whole population. Coexistence is promoted by higher mean path length, which leads to graphs with more segregated transmission clusters, but hindered by global and local efficiency. The more interconnected a network is, the harder it is for multiple strains to coexist. The reason why the clustering coefficient always acts against resistance is not entirely clear, but its interpretation is complicated by the fact that the treatment-clustering properties are partially determined by the level of clustering in the graph.

### Robustness of results to mechanism of treatment action

Throughout this paper, we have modeled the effect of antibiotic treatment as reducing the ability of an infected individual to spread infection to others. Hence, treated individuals are still infected, and therefore immune to infection with the other strain, but do not contribute to transmission (or contribute less, if treatment is imperfect, *ε* < 1). Alternatively, treatment could instead accelerate the rate at which an individual recovers from infection, a mechanism that has been used in many previous models of antibiotic resistance^30,31^. To determine if the mechanism of treatment action had any influence on our findings about the spatiotemporal patterns of resistance and to facilitate comparison of our results with those of other models, we considered a variant of the model where in treated individuals, the recovery rate increases from *g* → *g* + *τ* (see **Supplementary Methods** for details). We found that all of the results reported in the main text are recapitulated when treatment increases recovery rate, including the absence of coexistence in partially-treated well-mixed populations (Fig. S7C), a limited though larger parameter regime of coexistence in two-deme populations (Fig. S7E), common coexistence in multi-deme populations (Fig. S8B/C) and divergent resistance levels between neighboring demes receiving the same treatment level (Fig. S8E).

### Robustness of results to variations in treatment distribution and community structure

The results presented so far consider communities of demes where each deme is connected to the same number of other demes (e.g. “uniform networks”), where each deme is of the same size, and where there is either a 0% or 100% chance of receiving treatment if infected. While these simplifications allowed us to evaluate whether population structure alone could recreate observed spatiotemporal resistance trends in a minimal model, it is obviously not capturing many sources of variability that occur in the real world. Therefore, we also considered how relaxing each of these assumptions impacted our results. Populations described by graphs with heterogeneous connectivity (mean degree = 3, variance up to 10, Figure S9) supported similar levels of coexistence and between-neighbor differences in resistance levels as homogeneous graphs. Variation in deme size also did not seem to change the overall results (coefficient of variation up to 30%, Figure S10). We also created networks in which every deme was connected to every other deme at a fractional weight relative to the main network connections (Figure S11). Only as this level approached 1 did the likelihood of coexistence shrink. Intermediate levels of this background mixing (e.g. 0.01, 0.1) actually promoted robust coexistence (Fig. S11C), by allowing wild type and resistant strains to distribute throughout the population while still maintaining high and low treatment niches that selected for one or the other. Finally, we considered continuous distributions of treatment levels across demes, ranging from bimodal distributions where low and high levels of treatment were more common than intermediate levels, and unimodal distributions where the average treatment level was most likely and there was only a small variation between demes (Fig. S12). In general, extreme differences in treatment levels support coexistence over a wider range of parameters (Figs. S13, S14), but are not strictly necessary for it to occur and be distributed across most demes in the population (Fig. S15). When differences in antibiotic consumption across groups is small, coexistence only occurs when the average population-level drug efficacy is roughly equal to the cost of resistance (*ρε* ~ *c*, assuming groups are otherwise equal). In this case, other mechanisms may be needed to explain the pervasiveness of coexistence in the real world.

## Discussion

The prevalence of antibiotic-resistant bacterial infections displays several surprising spatial and temporal trends that defy canonical predictions of infection dynamics. The goal of this paper was to examine a simple and flexible model for competition between drug-sensitive and drug-resistant strains of an infection in a structured host population to see whether it could capture several of these trends. The first perplexing pattern observed in the data is the long-term coexistence of both sensitive and resistant strains at intermediate levels, which many other studies have attempted to explain^26,30–33^. Our simulations reproduce coexistence for a wide range of parameter values and suggest that it is a natural feature to be expected in spatially (or otherwise) heterogeneous populations with competing infection. In addition, we also captured more nuanced trends observed in surveillance data that other models have ignored. Regions that administer similar amounts of a particular antibiotic may experience persistent differences in drug resistance levels for many years, and these “frozen gradients” are observed even if regions are neighboring one another. These phenomena were reproduced even in our simple model (Figure 5A), but also occurred for more realistically variable population structures and treatment distributions (Figures S9–S15). We found that in a collection of partially-mixing subpopulations, subtle details about the pattern of connectivity and how treatment is distributed between them can lead to large differences in resistance prevalence between regions that otherwise seem similar (Figure 5A). We were able to identify specific covariates that contribute to resistance levels and to promoting coexistence with drug-sensitive strains. Similar to the use of “risk-mapping” to predict the distribution of vector-borne diseases based on environmental^55–58^ and human^58–62^ factors, these findings suggest that spatial risk-based assessment may be useful for the study of drug resistance. Surveillance on a finer scale will be needed to test these ideas on empirical data.

**Figure 5.**
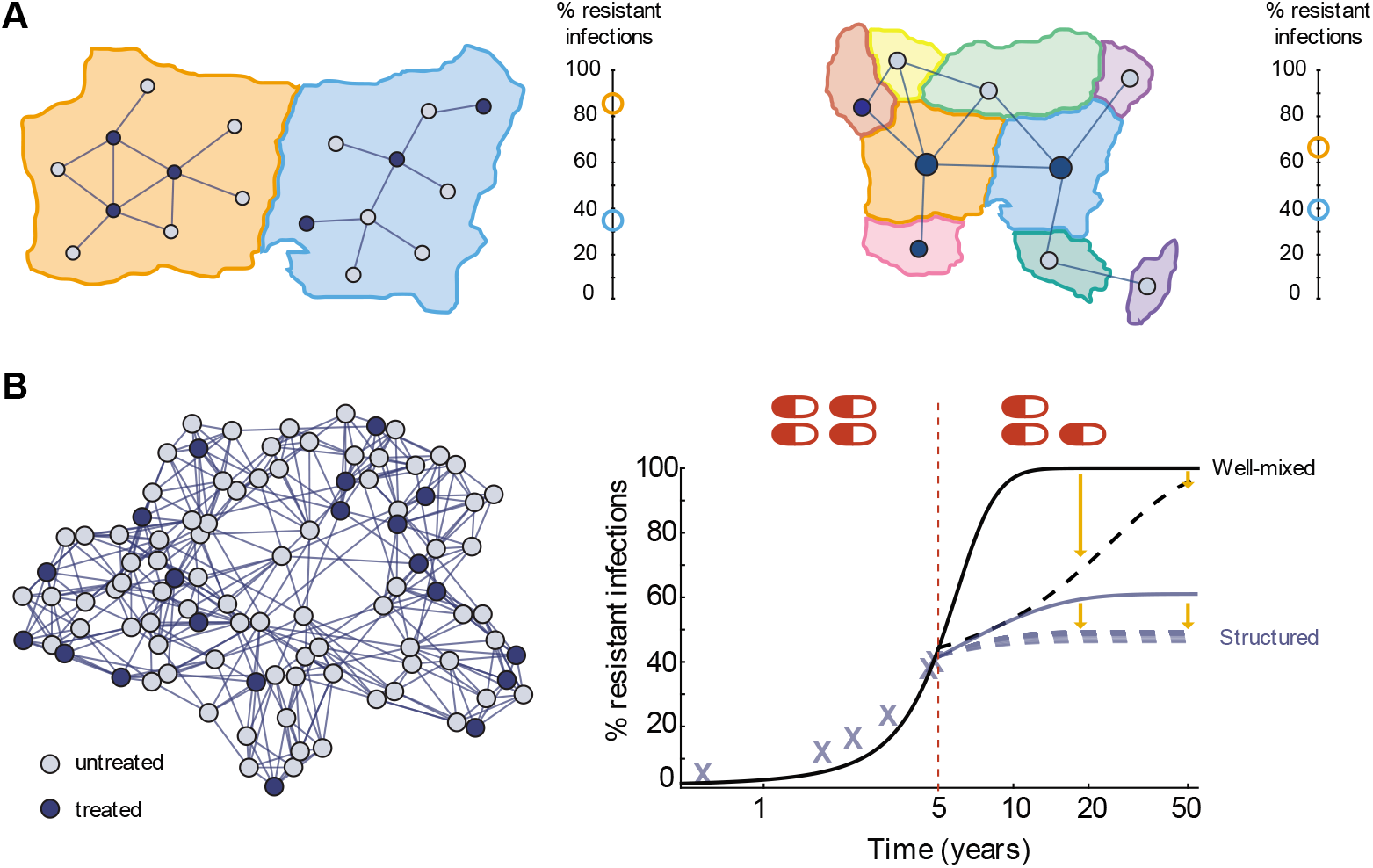
Implications for interpreting and predicting resistance levels. **A)** Our analysis suggests two possible explanations for large differences in resistance levels between two regions (orange and blue) receiving similar amounts of antibiotics. **Left:** The regions may differ in the distribution of antibiotic consumption within the population and its connection with the underlying transmission network. **Right:** The regions may differ in their connectivity to other regions which consume different amounts of antibiotics. **B)** Predictions about the impact of interventions on future resistance levels can be incorrect if population structure isn’t accounted for. **Left:** A population consisting of 100 sub-populations, with 20% treatment coverage at baseline. **Right:** Simulations of resistance levels in the “true” structured population (blue) with and without a hypothetical intervention (reduction to 15% treatment) applied at year 5. For comparison, predictions of a well-mixed model that with the same parameters (black), which approximates the dynamics before year 5 but diverges afterwards. The well-mixed model would predict large reductions in resistance in the short term but eventual fixation of resistance nonetheless (black dashed line), whereas the structured model predicts modest but sustained reductions which depend on the particular sub-populations targeted (blue dotted lines), and long-term coexistence. All simulations used kinetic parameters *κ* = 0.25/day, *β* = 0.05/day, *g* = 0.1/day, and *c* = 0.2. For the left of (A) and for (B), *ε* = 0.9, and for the right of (A), *ε* = 0.5. The population in (B) is a Watts-Strogatz network with degree 4 and re-wiring probability 0.1.

Increasing levels of antibiotic resistance are widely considered to be a major public health threat, and attempts are continually underway to forecast resistance levels with or without additional interventions using at least some type of mathematical modeling. Our findings highlight the fact that infection models which rely on assumptions of well-mixed populations are unlikely to be a useful tool for this task. As these models cannot reproduce ubiquitous trends observed in existing data on drug resistance epidemiology, they are unlikely to make predictions that are even qualitatively trustworthy (Figure 5B). For example, it is very important to know whether resistance for specific bug-drug combinations will tend towards 100% or settle at an intermediate level, and so any model must be able to explain the phenomena of coexistence for most strains. While clearly any degree of resistance complicates clinical management, the economic impact of partial vs complete resistance in a population could differ dramatically. In addition, by ignoring the details of spatial or demographic heterogeneity, we may miss an opportunity to more effectively target intervention or surveillance to particular subpopulations. The obvious difficulty is that such models may have to be highly-tailored towards a particular disease, and currently, there is a stark lack of data required to construct or calibrate realistic spatial models. While good-quality data is available on antibiotic consumption and drug resistance at the level of US states^63^ and European countries^14,16^, and could be combined with other datasets tracking inter-region travel, this is likely too large of a spatial scale to capture the relevant heterogeneity for this mechanism. More routine high-resolution mapping of resistance levels over space and time^39,64^, combined with spatial or demographic patterns of antibiotic consumption^45,65–68^ and data to estimate population connectivity patterns over which disease spreads, are sorely needed.

Throughout this paper we have considered a relatively simple type of population structure, while in reality it could be much more complicated. There may be multiple levels of structure - from a household to countries - with much more complicated patterns of connectivity and a continuum of transmission rates. Structures may be dynamic, as movement and interaction patterns of humans, animals, or disease-carrying material change over time. The mechanism leading to population structure varies depending on the infection of interest. For hospital-acquired infections, the movement of medical staff or the spatial arrangement of patients may be most important, while for community-acquired infections, the nature of close household, school, or workplace contacts may be of primary interest. For sexually-transmitted infections, networks of sexual contacts determine disease spread, while for diseases of livestock, relatively well-mixed contact within crowded farms along with patterns of transfer between farms may create an overall “meta-population” structure. Host population structures are notoriously difficult to measure in practice, but some approximation of them is likely needed to create realistic models for the spread of drug resistant infections. Technological advancements that allow better tracking of human mobility, such as wearable proximity censors^69–71^ or fitness trackers, cell phone location data^72–75^, or large-scale transit networks^76,77^, provide one source of population structure data. Advances in genetic epidemiology allow us to retrospectively trace the spread of infection between individual hosts or communities using pathogen phylogenies^78–82^, which also provides information on population structure. Only with better empirical characterization of transmission-relevant population structures and variations in antibiotic use can we test whether the mechanism described here can explain coexistence for specific bug-drug pairs.

Our model of competition between drug-sensitive and resistant strains is deterministic; we do not consider the stochastic early-stage invasion of an initial resistant mutant or the possibility of extinction of either strain. This approach is justified as we are interested in the dynamics of resistant strains that are already well-established and at high prevalence in the population. Since most antibiotics are closely related to natural products, resistance genes also naturally occur in environmental microbes, making “extinction” of resistant strains unlikely. Previous work by our group and others has examined how spatial heterogeneity or transmission network structure can accelerate the initial probability of emergence of drug resistance mutations^83–89^ and also on the loss of resistant strains when drug is withdrawn^90^. Models for the emergence of methicillin-resistant Staphylococcus aureus (MRSA) in which demes represent hospital vs community settings have found that stochastic effects can create a new regime of dynamics, whereby sporadic subcritical outbreaks can sometimes overwhelm control resources and push the system to endemic infection^91^. Even in our context of established resistance, some important differences could occur in a stochastic version of our model. If deme sizes are relatively small, the balanced coexistence of sensitive and resistant strains we observe at equilibrium could actually be a dynamic process of repeated colonization and extinction at the deme level (as described by metapopulation theory^92^). Extinction times for competing epidemics in homogeneous populations have only recently been estimated for homogeneous populations^93^, and the added complexity of our model suggests we are far from an analytic theory of invasion and extinction for infections in structured populations.

Our paper focuses on explaining trends observed for antibiotic resistance, but drug resistance is also a problem for antimicrobial therapy more broadly, including antivirals and antiparasitic drugs. To our knowledge, long-term coexistence of resistant and sensitive strains despite constant drug exposure has not been observed for HIV or malaria, the viral and parasitic infections for which targeted therapy is most widespread. In the case of HIV, rapid within-host evolution, including both *de novo* generation of resistant strains and reversion to susceptible strain in the absence of treatment would be expected to promote coexistence even in the absence of population structure. However, the prevalence of resistance appears to be changing rapidly along with dramatic changes in antiretroviral treatment availability and quality^94^. For example, the prevalence of resistance at the time of diagnosis (before treatment administration) has been increasing rapidly in South Africa and other low and middle income countries^95,96^, where widespread access to treatment is recent but follow-up care and resistance testing are still limited. In contrast, this primary resistance has been steadily decreasing in Europe, where treatment has been common since the 1990s albeit originally with suboptimal drugs (e.g.^97^). For malaria, it appears that resistance levels in a population continually increase under drug exposure, as extremely high levels of chloroquine or sulfadoxine/pyrimethamine resistance have been observed (e.g.^98^). However, antimalarial treatment is not generally individualized and international guidelines suggest that country or region-wide treatment recommendations switch to new drug classes once suspected resistance is above a threshold of around 15-25%. Consequently, the removal of selection for those resistance strains and their subsequent decline(e.g.^99^) likely precludes the occurrence of stable coexistence. Interestingly, a recent modeling study examining the role of co-infection with resistant and sensitive strains of malaria showed how this can suppress selection for resistance^100^, very similar to recent ideas suggested for promoting coexistence in commensal bacterial infections^32^.

Spatial heterogeneity has been understood by population geneticists to be an important factor in the spread of genes since the days of Fisher^101^ and later Kimura^102^. Other classic work uncovered mechanisms by which the coexistence — as opposed to competitive exclusion — of species could be promoted by spatial heterogeneity^103,104^. More recently, Gavrilets & Gibson^105^ examined a two-deme system with two competing types, each of which has higher fitness than the other in one of the two demes, and observed a phase diagram very similar to that for our two-deme infection model (Fig. 3), in which polymorphic equilibria were possible. This idea was later extended to multiple connected demes^106^, where it was demonstrated that spatial heterogeneity can preserve a species in spite of fitness differences which in a well-mixed model would drive it to extinction. Other work involving more complex population structures^107^ or more complex fitness distributions across space^108,109^ has examined how the rate of invasion of rare mutants depends on spatial heterogeneity. This previous work all involved traditional population-genetic models, such as Moran or Wright-Fisher processes, and before this study it was not clear which results would generalize to infection dynamics models. For example, the role of network structure in modulating the fixation probability of new strains differs dramatically between Moran processes^107^ and infection models^88^.

Several other recent studies have examined the role of population structure in facilitating coexistence of drug resistant infections. Kouyos et al.^110^ considered populations divided into community members and hospitalized patients, and examined if such structure could support the coexistence of two separate strains of methicillin-resistant *Staphylococcus aureus* (MRSA). Similar to our model one strain was more resistant yet more costly, and was favored in more highly-treated hospitalized individuals, while the other strain was less resistant but more transmittable. They found a relatively narrow parameter regime supporting coexistence, which could be increased only somewhat if they additionally structured the population into different age groups. Interestingly, they found that beyond the parameter region that supported coexistence at equilibrium, there was a much wider regime leading to decades-long transient coexistence, suggesting that our estimates of the extent of coexistence are likely very conservative underestimates. Cobey et al.^31^ examined the role of age structure in supporting coexistence of drug sensitive and resistant strains of *S. pneumoniae*, and found that this mechanism could lead to coexistence but only for a relatively narrow range of costs of resistance. In addition, their model included a mixture of different serotypes, and they found that apparent population-level coexistence often occurred due to summing many unique serotypes which were each nearly all resistant or all sensitive. Similar to us, Blanquart et al.^33^ considered a population consisting of a set of interconnected “demes” with different treatment levels, but in contrast, for mathematical tractability, they restricted their analysis to structures in which either all demes are connected to all others with equal weight, or are completely isolated. Their model was parameterized such that the maximal force of infection from all outside demes equaled to the force of infection from within the same deme, whereas in our model the force of infection from outside can be much larger than from inside. They also concluded that population structure can support coexistence, but in contrast to our findings, under their model assumptions coexistence only occurs with very weak inter-deme mixing. By considering larger and more variable transmission network structures and treatment distributions, our work thus suggests the potential for a more prominent role of population structure in supporting coexistence. Compared to previous work, our criteria for the occurrence of coexistence in models is more stringent, since we require at least 5 % prevalence of each strain (compared to >0%^33,110^ or >2%^31^), and additionally require coexistence within each individual subpopulation (“robust coexistence”). Our paper additionally differs from these previous works by going beyond coexistence to examine other common spatial trends in resistance data (i.e. trends **(II)** and **(III)**), and by comparing the role of a range of spatial metrics describing population structure and treatment allocation patterns in promoting coexistence (Table 1).

While our results show that population structure can explain observed spatiotemporal patterns of antibiotic resistance, it is unlikely to be the only mechanism contributing to either long-term coexistence or differences between regions. Persistence of both drug-sensitive and drug-resistant strains is also promoted by co-infection and superinfection^30–32^, by overlaps between resistance status and serotype^30,31^ or other traits under balancing selection^36^, and by heterogeneities in transmission or recovery rates as opposed to treatment coverage^30,31^. Regions receiving similar overall levels of drug but experiencing different frequencies of resistant infections could alternatively be explained by local differences in i) prescribed vs consumed doses, ii) who receives drugs (e.g. age group, hospital vs outpatient) and for what type of condition, iii) genetic background of the pathogen and the cost of resistance, iv) transmission or recovery rates, or v) strain-specific control measures. For example, take two regions with identical transmission networks, drug consumption, prescribing behavior, and infection burden. If one region institutes a control policy whereby individuals infected with the drug-resistant strain are isolated, then in that region the cost of resistance (*c*) is effectively higher - even for a genetically identical strain - and the resistance level will be lower. It is difficult to judge precisely how realistic our results are, since our models of infection dynamics and population structure are both dramatically simplified, but it likely cannot explain coexistence over the entire range of parameters for which it is observed empirically. In addition, in our results it is relatively rare to see large differences in resistance prevalence between regions with similar treatment rates unless they have very different connectivity patterns within or between them, and unless there are large spatial or demographic heterogeneities in treatment use that overlap with clustering of disease transmission. In reality, multiple mechanisms probably act in concert to explain the observed patterns. Moreover, there is unlikely to be a single explanation for each different bug-drug pair for which these trends are observed. Other disease-specific factors that may be relevant to the ecology of antibiotic resistance include whether the bacteria is primarily pathogenic or also a commensal colonizer, whether infection is more common in the community or in healthcare facilities, whether resistance is carried on plasmids or chromosomally, and whether there are environmental reservoirs and if they are exposed to antibiotics.

## Methods

### Model assumptions

Our two-strain model of infectious disease dynamics makes a number of simplifying assumptions. We assume that individuals can transmit the infection immediately upon becoming infected, hence ignoring any sort of “latent” phase or “exposed” class. Birth, death, and migration into and out of the population are not modeled. The infection is assumed to be non-lethal, and may even represent a non-pathogenic “colonized” state with a commensal organism. Infected individuals all recover to be susceptible again, which presumes that there is no long-lasting immunity to re-infection. We do not allow for any co-infection (simultaneous infection with both strains) or super-infection (replacement of one strain within an individual by another), which implies that there is complete cross-immunity during infection. We model resistance as a binary trait: there are only two strains, either completely sensitive to the drug, or completely resistant. In reality, there may be strains with intermediate levels of resistance.

Individuals who are assigned a status of “treated” will receive treatment immediately upon being infected, every time they are infected. This ignores the possibility that there may be a delay to receiving treatment or that treatment behavior may vary within an individual over time. Treatment is assumed to reduce transmission of the infection, but may not be perfectly effective at doing this (which could approximate the effect of treatment delay or imperfect drug efficacy). In the **Supplementary Methods**, we consider an alternative model in which treatment acts to increase the rate of recovery from infection instead. In reality, treatment likely has both effects, since it generally acts to reduce pathogen loads within a host, which leads to reduced contagiousness and reduced duration of infection. Although we track uninfected (susceptible) individuals as either “untreated” or “treated”, our results are independent of whether uninfected individuals actually ever receive drug.

Our population structure consists of *M* subpopulations (“demes”) of equal size *D* for a total population of size *N*. The structure is assumed to be static. For each model, the transmission rates *κ* and *β* were scaled so that the model behavior did not depend on *N*, *D*, or *M*. In the one- and two-deme case, *κ* was multiplied by *N*, and in the multi-deme cases, both *κ* and *β* were multiplied by *D*. A more detailed derivation of this scaling is given in the **Supplementary Methods**. This scaling is equivalent to saying that individuals are limited in how many contacts they can have with others, and that this limit is independent of the total population size.

By using differential equations to model infection, we assume that the population sizes of individual demes are large enough so that variation from the average behavior is not important. We additionally assume that resistance is pre-existing and that stochastic extinction of either strain from the population is not possible. For the most part, we examine only the equilibrium behavior of the models.

### Model equations

#### Well-mixed population (single deme)

Infection spreads between all individuals with the within-deme transmission rate *κ*. A fraction *f* of all individuals are treated with drug immediately upon infection by either strain. We track the proportion of the total population who are infected with the wild type and untreated (*w_u_*), infected with the wild type and treated (*w_t_*), infected with the resistant strain and untreated (*r_u_*), and infected with the resistant strain and treated (*r_t_*). The number of uninfected (susceptible) individuals is given by *s* = 1 – *w_u_* – *w_t_* – *r_u_* – *r_t_*. The dynamics are then described by the following set of differential equations (see **Supporting Information** for a derivation):

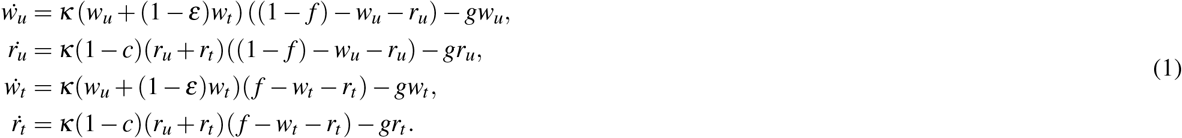

#### Two deme population

Infection spreads between all individuals in the same deme with rate *κ*, and between any two individuals in different demes at rate *β*. The demes are of equal size *D*. Treatment is assigned to all individuals within only one of the two demes. We track the proportion of individuals in the treated deme that are infected with the wild-type/drug-sensitive strain (*w_u_*) and infected with the resistant strain (*r_u_*), and the same proportions for the treated deme (*w_t_* and *r_t_*). The number of uninfected (susceptible) individuals in the untreated deme is *s_u_* = 1 – *w_u_* – *r_u_* and in the treated deme is *s_t_* = 1 – *w_t_* – *r_t_*. The system is then described by the following set of differential equations (see **Supporting Information** for a derivation and generalization to unequal deme sizes):

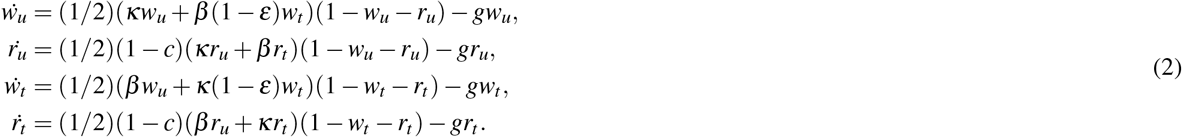

#### Multi-deme population

Infection spreads between all individuals in the same deme with rate *κ*, and between any two individuals in different but connected demes at rate *β*. There are *M* demes each of size (*D*). Each deme may only be able to spread infection to a sub-set of other demes, and the connectivity of the population is described by the adjacency matrix Δ*_ij_* (Δ*_ij_* = 1 if an individual in deme *j* can be infected by an individual in deme *i*). The *degree* of deme *i*, *d_i_*, is the number of neighbors it is connected to, *d_i_* = ∑_*i*_Δ_*ij*_. Treatment is assigned at the level of the deme, described by indicator variable *T_i_* (where *T_i_* = 1 if deme *i* is drug-treated and *T_i_* = 0 otherwise). The fraction of demes that are treated is *ρ*. The system of equations describing the fraction of individuals in each deme who are infected, with either the wild-type (*w_i_*) or drug-resistant strain (*r_i_*) is:

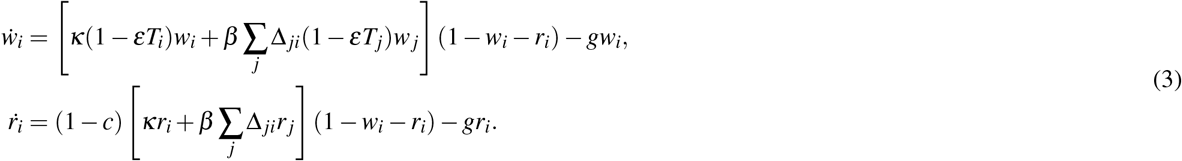

A derivation that is generalized to the case of demes of unequal size and with arbitrary treatment levels in each deme is provided in the **Supporting Information**. Note that in constructing the adjacency matrices Δ*_ij_* we ensure that the graph contains a single giant component, and not multiple isolated parts.

Derivations for the equilibria, stability conditions, and basic reproductive ratio (using the next-generation technique^111^) are given in the **Supplementary Information**.

### Numerical results

The differential equations were numerically integrated in MATLAB, using the Runge-Kutta solver provided in the function `ode45`. The initial condition for each parameter-population structure combination was each deme having a level of infection with the wild type that would occur if there were no resistant strain (a fraction 1 – 1/*R*_0_ of individuals are infected with the wild type), and a very small level of infection with the resistant strain (10^−3^). The rest of the population was uninfected.

The system was integrated until an equilibrium was reached, which was defined as the point at which the sum of the time derivatives for the fraction of each strains in all demes became less than 10^−4^/*M* per day, where *M* is the number of demes. To ensure that the equilibrium reached by the solver is stable, once an equilibrium was reached by the above standard, we applied a small random fluctuation with magnitude not exceeding 1% to the value of all strains in all demes. If the same equilibrium was arrived at after repeating this process three times, the equilibrium was taken to be sufficiently stable and used as the result of that simulation. We also explored using more stringent thresholds, but found that our results were nearly always within 0.01% of the value that would be achieved and reduced simulation times (Supplemental Figure S16). To determine whether there was only a single positive and stable equilibrium for the multi-deme system for a given structure and parameter set, we simulated the same systems many times with uniformly-at-random initial conditions. For each deme, the initial fractions [*w_i_*, *r_i_*] were both drawn from the uniform distribution on [0,0.7]. If their sum exceeded 1, both were re-drawn. Previous work has proven mathematically that stable equilibria exist for these systems^112,113^. We found that the same equilibrium was always reached (Figure S17). More details are given in the **Supplementary Information**.

### Network generation

#### Random regular graphs

Random regular graphs are randomly-generated graphs in which every node has the same degree (number of incident edges). There are multiple existing algorithms to create random regular graphs. Our random regular graphs were generated according to a pairwise-construction mechanism^114^ implemented in MATLAB^115^. For most results presented in the paper we used networks with *M* = 20 total nodes and degree *d* = 3.

#### Watts-Strogatz graphs

The Watts-Strogatz algorithm is a method of constructing networks with *M* total nodes and *M* * *d* total edges, but with heterogeneous properties. In the original construction, a ring lattice is formed with each of the *M* nodes connected with *d* edges to neighbors in a ring formation. Then, with probability *p*, the target of each edge is rewired to a uniformly random node. For our 1000 graphs, we selected *M* = 50, but allowed degree *d* to be selected uniformly-at-random for each graph from {3, 4, 5, or 6}, and allowed *p* to be selected uniformly-at-random for each graph from [0.1,0.6]. The resulting graphs have a broad distribution all graph properties of interest (Figure S3) and of the frequency of resistance (Figure S4).

### Network statistics

A set of graph-theoretical properties were tested for their relationship to resistance levels within demes and within populations. A description of these properties is given below.

#### Eigencentrality

For a node *i*, its eigencentrality is a measure of not only its centrality (having a high degree, for example), but is also weighted according to how high the centrality scores of its neighbors are; as an example, an academic paper with many citations may have high centrality, but if the papers citing it have themselves low centrality then the eigencentrality of the original paper may be low. Formally, the eigencentrality is the leading right eigenvalue of the adjacency matrix, and so satisfies the implicity equation *E_i_*: *E_i_* = (1/*λ*) ∑_*j*_Δ_*ij*_*E_j_*, where *λ* is a normalization factor.

#### Global efficiency

In general, the average efficiency of a graph is defined as 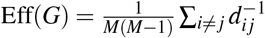, where *M* is the number of demes/nodes and *d_ij_* is the smallest number of edges between nodes *i* and *j*. Therefore, graphs that have higher efficiency have on average shorter path lengths between nodes. Since *M* = 50 is fixed in all of our structures examined in this section, the only varying factor is the distances between nodes; the smaller the distances between all given nodes, the higher the efficiency. Global efficiency involves comparing the graph to the most efficient possible graph with as many nodes. Since this simply involves another prefactor which is equal for all of our graphs, our global efficiency is proportional to the average efficiency.

#### Local efficiency

Local efficiency is defined as Eff_*l*_(*G*) = *M*^−1^ ∑_*i*_Eff(*G_i_*), where *G_i_* is the subgraph in which we keep the neighbors of node *i* but remove *i* itself, and the efficiency is defined as above.

#### Clustering coefficient

The network average clustering coefficient was calculated by averaging the local clustering coefficient over all nodes. The local clustering coefficient is calculated for a node *i* as the fraction of all possible edges that could exist between the neighbors of *i* which actually do exist in the network. Said another way, the local clustering coefficient is determined by counting all triplets (sets of three connected nodes) centered on node *i* and then calculating the fraction of these triplets that are triangles (all three nodes connected to each other).

#### Mean path length

Given any two nodes, we can calculate the path length as the smallest number of edges that must be traversed to move from one to the other. The average path length is the average over the distribution that arises from calculating this smallest number for all possible pairs of nodes.

### LASSO regression

LASSO (least absolute shrinkage and selection operator)^116^ is an advanced regression technique that favors sparse models. This is accomplished by forcing the sum of all the regression coefficients to be less than a certain number; in optimal fitting, some coefficients are then set to zero (i.e., deemed unimportant) rather than assigned a non-zero value. This is usually accomplished by adding the sum of all the regression coefficients, multiplied by a regularization parameter *λ*, to the quantity being minimized (such as sum of the squared error). When *λ* is large, having few covariates in a model becomes more important than the model fitting the data accurately. Therefore, the error-minimizing model usually occurs for some intermediate value of *λ*.

We employed Matlab’s `lasso` (and `lassoPlot`) function, which imposes the LASSO constraint on the **L**_1_ norm of all fit coefficients onto a linear regression. In the ”deme-level” regression, the outcome variable was the proportion of infections that were with the resistant strain within that deme, and the possible predictors were local properties of that deme. In the ”population-level” regression, the outcome variable was the proportion of resistant infections across the entire population, and the possible predictors were properties of the entire population graph. At each level, the regression was conducted both under low resistance level conditions (24% demes treated, leads to < 50% resistant) and high resistance level conditions (40% demes treated, leads to > 50% resistant).

Although a rigorous and robust significance test for the LASSO is still lacking, we can compute the mean-squared error of the predictions of the model to the actual data for cross-validation. Here we choose five-fold validation, meaning that the data is randomly split into five chunks of equal size. Four of these chunks are used to fit the regression, and then the mean-squared error between the actual fifth chunk and its prediction by the regression is measured. If this mean-squared error is sufficiently small, the model is a good fit. The results of this cross-validation procedure for LASSO regression fits to both deme and population-level properties are shown in Fig. S5.

The rank of each property displayed in Table 1 is determined from the magnitude of the regression coefficient at the value of the regularization constraint *λ* that minimizes the mean-squared error after five-fold cross-validation (Fig. S5, S6), averaged over both the low and high treatment case.

## Acknowledgments

We thank Jeff Gerold, Sam Sinai, Kamran Kaveh, Anjalika Nande, Gabriel Leventhal, and participants in the 2018 Gordon Research Conference on Drug Resistance for helpful discussions.

## Funding

This work was supported by the National Institutes of Health (DP5OD019851) and the Bill & Melinda Gates Foundation (OPPOPP1148627).

## Author Contributions

M.S.K and A.L.H. conceived the study. M.S.K., M. A. N, and A.L.H. analyzed the model. M.S.K. performed all simulations, network analysis, regression, and stability analysis of the model and analyzed the resistance data. T.L.A. and C.A.D. derived and analyzed the model for continuous treatment and heterogeneous deme sizes. M.S.K., M. A. N, and A.L.H. interpreted the results and wrote the manuscript.

## Competing interests

The authors declare that they have no competing interests.

## Data availability

All data shown is available from the original source at the links provided in the references. All equations and algorithms needed to generate the results of this paper are described in the Methods/Supplementary Methods, and code to simulate the model equations is available from the authors by request.

## Supporting Information

### Analytic results for well-mixed single deme with treatment

The dynamics of competition between drug-sensitive and drug-resistant strains of infection in an infinite, well-mixed population where a fraction *f* of individuals receive treatment if infected is described by Eqns. 1. This system of equations tracks the *proportion* of the population in each treatment and infection category (infected with the wild type and untreated (*w_u_*), infected with the wild type and treated (*w_t_*), infected with the resistant strain and untreated (*r_u_*), and infected with the resistant strain and treated (*r_t_*)). These equations were developed by first writing down the dynamics of the *number* of individuals in each state, including uninfected (susceptible):

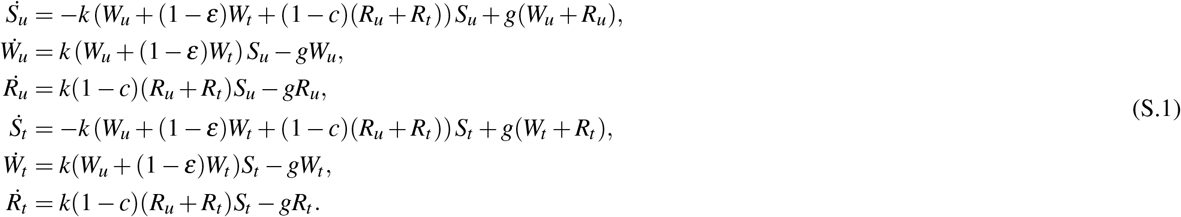

The transmission rate *k* represents the rate at which each infected individual contacts and infects each susceptible individual. Although we use the terms *S_t_* and *S_u_* to track susceptible individuals who are “treated” or “untreated”, our model is agnostic as to whether these uninfected individuals actually receive drug, or if these markers simply indicate *if* these individuals will (or will not) receive treatment *when infected*. While here we assume a pre-determined fraction *f* of the total population will receive treatment if infected, similar models could be constructed assuming the decision to receive treatment is made instantaneously upon infection with probability *f*, or that there is some rate of receiving treatment, such that a fraction *f* will receive treatment before recovery. The analysis of these models gives identical steady state results.

Because in our model the total population size *N*, as well as the fraction of individuals who will (or will not) receive treatment if infected, are both constant, we can remove the variables for susceptible individuals using *S_u_ = N*(1 – *f*) – *W_u_ – R_u_* and *S_t_ = N f – W_t_ − R_t_*. To further reduce the variables in this system and remove any dependence on total population size, we let *κ = kN* and *x_i_ = X_i_/N* (the fraction of individuals instead of the total number, for *X = W,R* and *i = u, t*), which then leads to Eqns. 1. By keeping *κ = kN* as the constant parameter, we are implicitly assuming that the total rate at which a single individual can contact and infect others when the population is completely susceptible (*kN*) is independent of the population size. This is a reasonable assumption for many infections which require some type of active contact, and here allows us to reduce the number of parameters required. None of our results depend on this scaling, and the effect of changing population size is equivalent to changing *κ* in this deterministic model.

There are three possible equilibria of Eqns. (1). In each of these equilibria, there is no possible coexistence of infections. These equilibria are as follows:

1. **No infection survives**

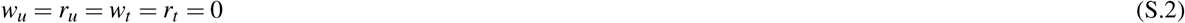 This uninfected equilibrium is only stable when

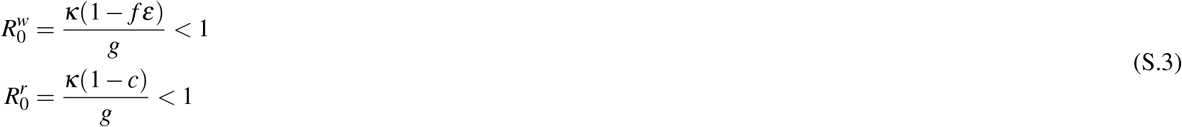
2. **Only wild-type (drug-sensitive) strain survives**

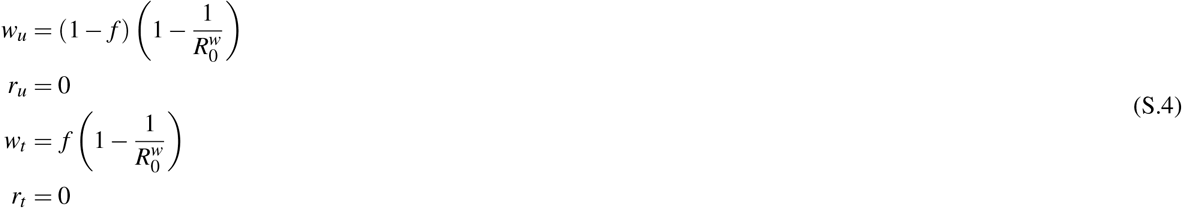 This infected equilibrium is stable when

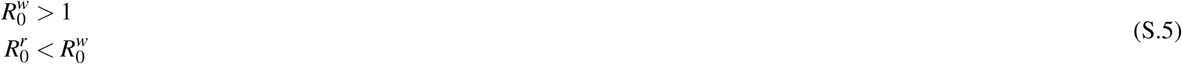

where the later condition implies *c > fε*.
3. **Only drug-resistant strain survives**

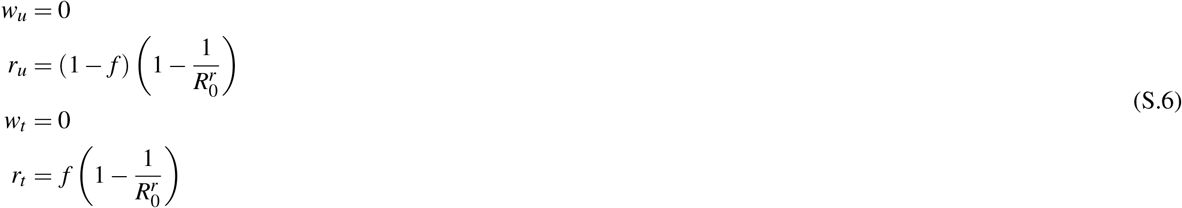 This infected equilibrium is stable when

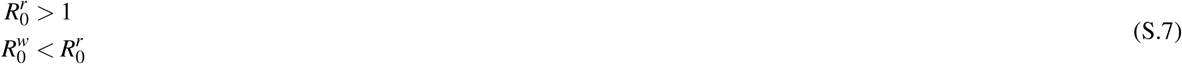

where the former condition implies *c < f ε*.

### Analytic results for two connected subpopulations

In an effort to be more general than in the main text, we consider the case of a population of total size *N* divided into two demes, with a fraction *f* of the total population is contained in the drug-treated deme. The system describing the *number* of individuals in each state can then be written as:

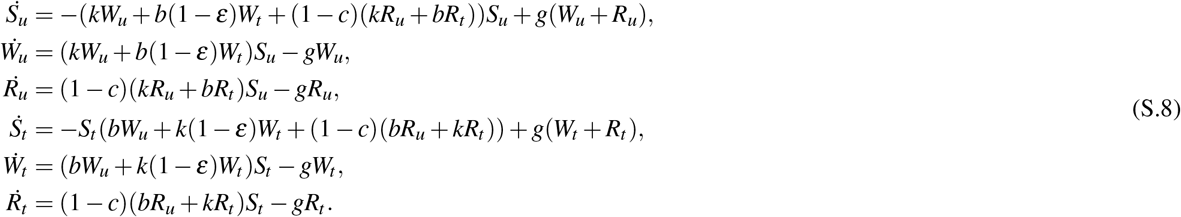

Since the size of each deme is fixed, we can remove the variables for susceptible individuals using *S_u_* = *N*(1 – *f*) – *W_u_ – R_u_* and *S_t_ = N f – W_t_ – R_t_*. To create a nondimensional version of the system, we instead track the *fraction* of individuals in each deme who have a particular infection status, so *w_u_ = W_u_*/(*N* (1 – *f*)), *R_u_ = R_u_*/(*N*(1 – *f*)), *w_t_ = W_t_*/(*N f*), *R_t_ = R_t_*/(*N f*), and *κ* = *kN* and *β = bN*, which gives the altered system:

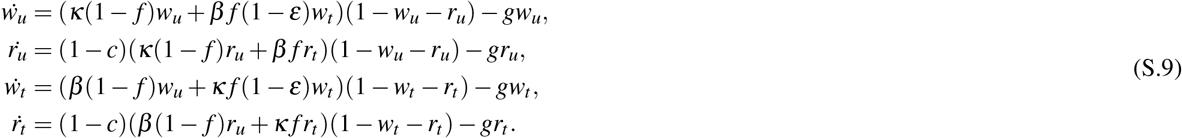

Using the next-generation technique^111^, we arrive at the following values for the basic reproductive ratios,

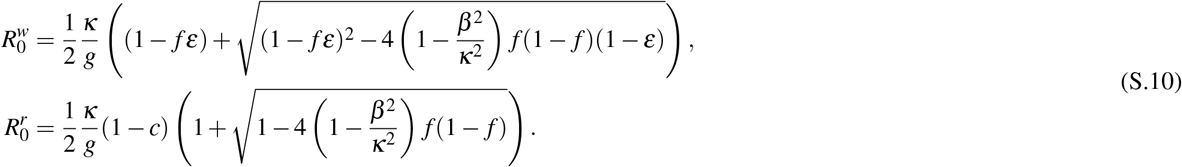

Equations (S.9) and (S.10) reduce to Eq. (2) and

In general, this system has nine equilibria. However, only one of these is stable and physically realizable (e.g. having between 0% and 100% of individuals infected) for any given parameter set. These are the values that are plotted in the main text. These outcomes fall into one of two qualitative categories. Either one strain drives the other to competitive extinction (true in Fig. 3 for *c* > 0.4 and *c* < 0.06, for instance) or both strains exist in both demes simultaneously. The boundaries separating this region from the competitive-exclusion region are 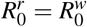, the lower bound when viewed as a function of the cost of resistance *c*, and the curve which solves 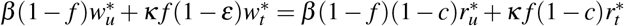, where 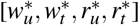 denote the equilibrium values. This second curve, in words, is where the drug-treated population is infected at the same rate by both drug-resistant and drug-sensitive strains.

For the parameters chosen here and used throughout the paper, it is impossible to have 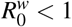. For sufficiently high *c*, we can have 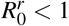. However, this line in parameter space lies deep within the region where the population is infected with the wild-type strain only.

### Derivation of a multi-deme model

Here we derive the general case for the spread of wild-type and resistant strains of infection in a population consisting of multiple connected demes (with connectivity described by adjacency matrix Δ), where demes may have different sizes (*D_i_*) and different treatment levels (*f_i_*). The dynamics of infection are described by the following system of equations

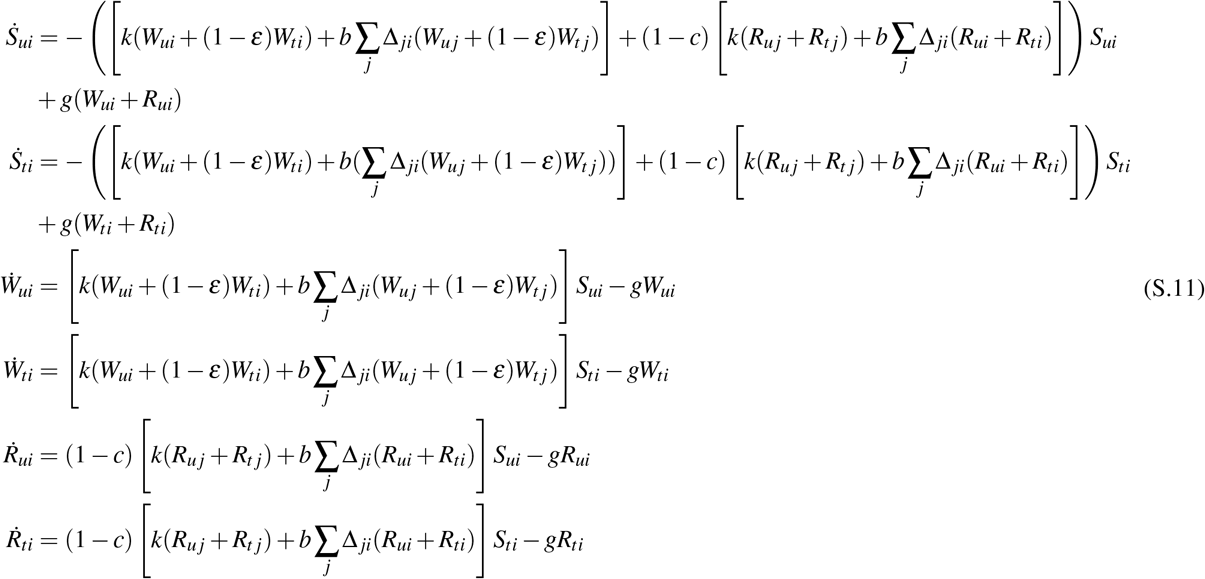

where *X_it_* represents the total number of individuals in deme *i* infected with strain *X* (or susceptible, *S*) who will be treated if infected, whereas *X_iu_* is those who won’t be treated if infected.

Since the population size *D_i_* of each deme is constant and we assume it is predetermined who will get treatment *if* infected, we can remove the equations for the susceptible individuals using *D_it_ = f_i_D_i_ = S_it_ + W_it_ + R_it_* and *D_iu_* = (1 – *f_i_*)*D_i_ = S_iu_ + W_iu_ + R_iu_*.

To derive a dimensionless form the the equations, we let *x_i_ = X_i_/D_i_*, and define 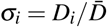 where 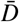 is the average deme size. We define 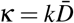 and 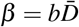, to derive the simplified and scaled equations

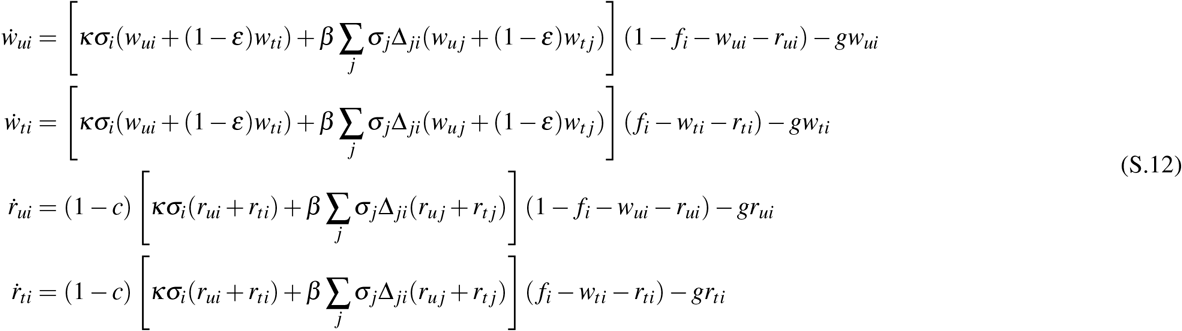

In the limit of either 0% or 100% treatment in a deme, this system reduces to

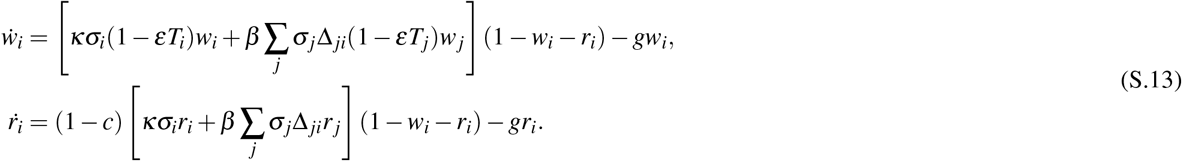

In the limit of both constant deme size and 0% or 100% treatment in a deme, this system reduces to Equation 3.

### The apparent absence of multiple equilibria

Although the generic system of equations modeled by Eqns. (3) admits multiple equilibria, we actually never observed any dynamic phenomena other than a steady convergence to a single stable fixed point, even when initial conditions were wildly different from that considered in the main text. It is not uncommon for infection models to admit equilibria which are physically unrealizable, i.e., taking values outside the unit simplex — it may therefore be the case that any other stable equilibria are simply unphysical.

However, to comprehensively demonstrate this fact — either theoretically or numerically — is outside the scope of the current work. To prove such a result theoretically would necessarily draw on ideas of nonlinear algebra or algebraic geometry, such as intersection theory, which are outside the present scope; to demonstrate numerically that for all parameter sets and graph structures there is only one stable equilibrium is computationally infeasible, even on graphs of a given small size.

To illustrate the idea that there seems to be only one stable equilibrium per system, we fixed an exact distribution of treatment with *ρ* = 0.35 to be used on every random regular graph used in the main text. This means that, since all nodes in all graphs are labelled (and the random regular graphs in the main text have twenty demes each), seven labelled demes (here 1, 3, 8, 11, 15, 19, 20) were drug-treated irrespective of graph structure. We then performed, for all 30 graphs, 100 trials with uniformly random initial conditions over every deme, i.e., every deme can start with any possible arrangement of wild-type and resistant infectives (so long as the sum of the two does not exceed 1). We then compared the equilibria seen with the equilibria found for the initial condition used in the main body of the text.

There are many possible metrics which would illustrate the relative closeness of all the equilibria seen. In Fig. S17 we have elected to plot the maximum matrix norm (the 2-norm) of the difference between the equilibrium seen in any of the 100 trials and the equilibrium from the main text. That is to say, all of the trial equilibria differ in norm from the main-text equilibrium by an amount not more than that plotted in the figure for a given graph structure. The parameters used were our default: *κ*=0.25, *β*=0.05/day, *g*=0.1/day, *ε*=0.9, *c*=0.2.

In all of these cases, the maximum norm deviation is extremely small. This could be due to two possible reasons. The first is due simply to aggregated numerical error from the many steps involved of a) calculating the equilibrium (which involves a threshold of stopping when the time-derivatives in all demes fall below a certain threshold, which can allow for some very small freedom in what an “equilibrium” is in a given trial), b) calculating the matrix norm (which amplifies these small errors). The second explanation is that while multiple stable equilibria may exist, they are extremely close together, with almost clinically-unmeasurable differences in the rates of wild-type and resistant infections within a given deme in a given community.

### Model with effects of treatment on recovery rates

To demonstrate the robustness of the results described in the main text, we explored a second model in which the drug, rather than inhibiting the spread of the infection, instead speeds up recovery. The effect of treatment is to increase the rate of recovery from *g* to *g* + *τ*. In this model formulation a perfectly efficacious drug would correspond to *τ* → ∞ (instantaneous clearance of infection). A finite value of *τ* takes into account the fact that it takes some time for an infected individual to seek a clinician, obtain a prescription for drug, and clear the infection as a result of treatment.

This model matches qualitatively with that discussed in the main text, in that in reproduces the spatial motifs in the same way. The two models are also quantitatively quite similar for reasonably-akin parameter sets. Here we reproduce the main results for this alternative model.

#### A single well-mixed deme

Here, the system can be written as

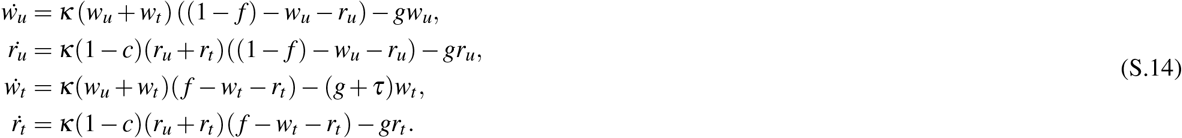

The basic reproductive ratio *R*_0_ for each strain alone is

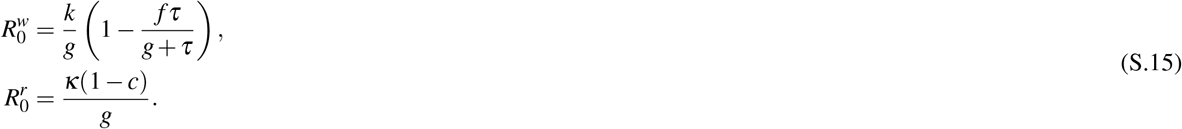

For the purpose of comparing this model, the value of *τ* for which 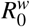 in the main model (with our baseline *ε* = 0.9) equals 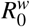 in this alternative model is *τ* = 0.9. This system admits only three stable, physical (having fractions of treated individuals between 0 and 1) solutions: one in which all infections die out 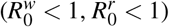, one in which only wild-type infections persist 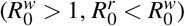 and one in which only resistant infection persists 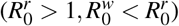. There is no possibility for coexistence. The condition for resistance to persist (instead of wild type)is 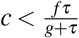.

#### Two connected demes

In this model, the equivalent of Eqs S.9 can be written as

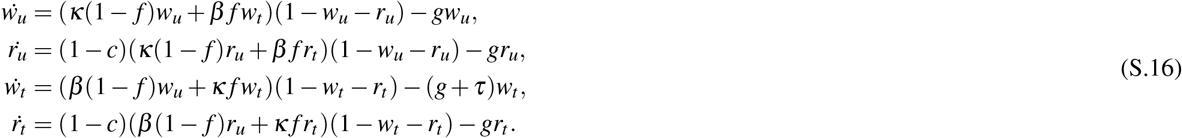

Using the next-generation technique, we were able to derive the values of *R*_0_ in a two-deme system for the alternative model, as an analog to Eq. S.10:

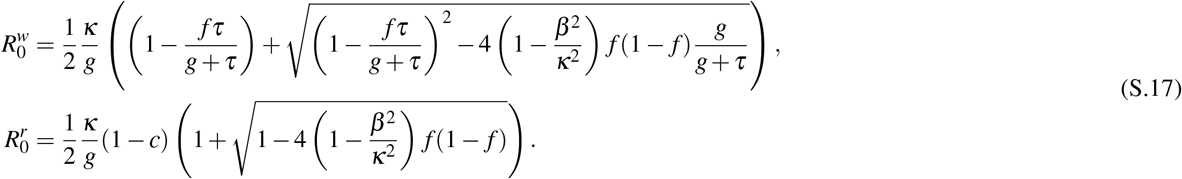

The expressions are slightly more complicated, but the curves generated are similar to the main text. Fig. S7 shows the analog of Fig. 3. As in the main text, here the boundaries of the coexistence region are the curves 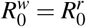 (lower boundary) and the implicit curve where more wild-type than resistant infected individuals are produced per unit time in the drug-treated deme (upper boundary).

#### Multi-deme population

The scaled version of the multi-deme model presented in Equation (3) for the case of treatment acting on the recovery rates is

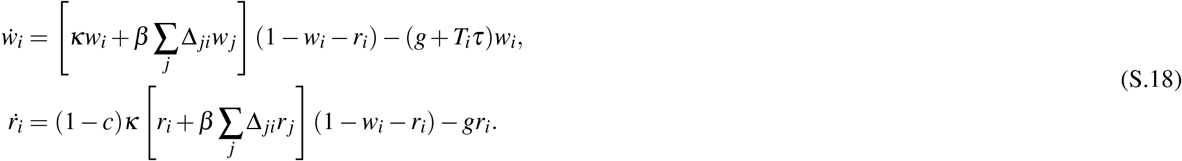

## Supplementary Figures

**Figure S1.**
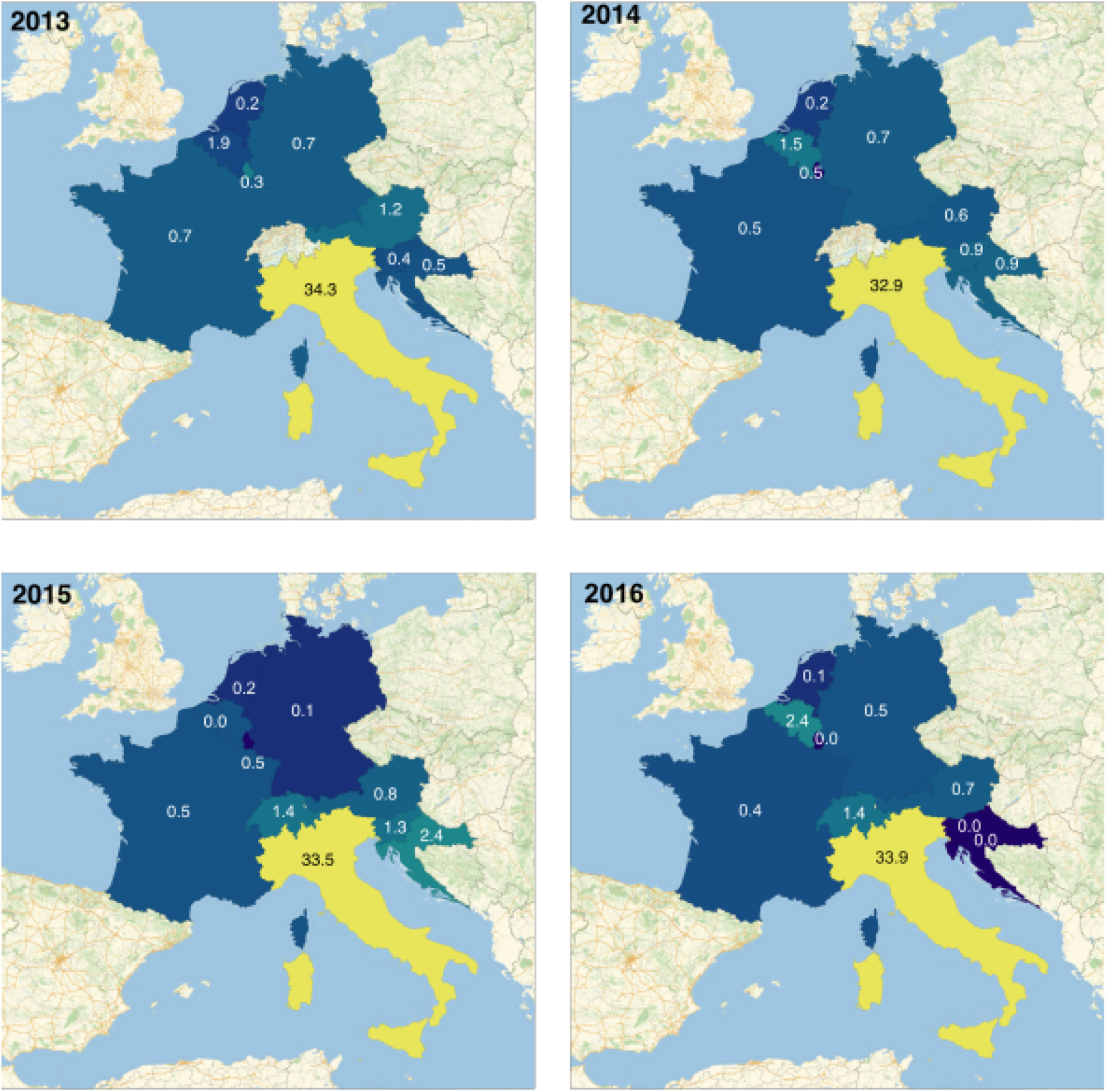
Stable spatial heterogeneity in resistance levels in Europe. Percent of *Klebsiella pneumoniae* isolates resistant to carbapenems in Austria, Belgium, Croatia, France, Germany, Italy, Luxembourg, and Slovenia from 2013-2016 as reported in the ECDC’s European Antibiotic Resistance Surveillance Network^13,14^ and Switzerland as reported by the Swiss Centre for Antibiotic Resistance from 2015-2016. Each country is labeled with the resistance level (%). The year-to-year deviation in Italy from the average value shown in Figure 1 is less than 1% for all years. For all other countries, the frequency of resistance never reaches even 1/10 of the average value seen in Italy.

**Figure S2.**
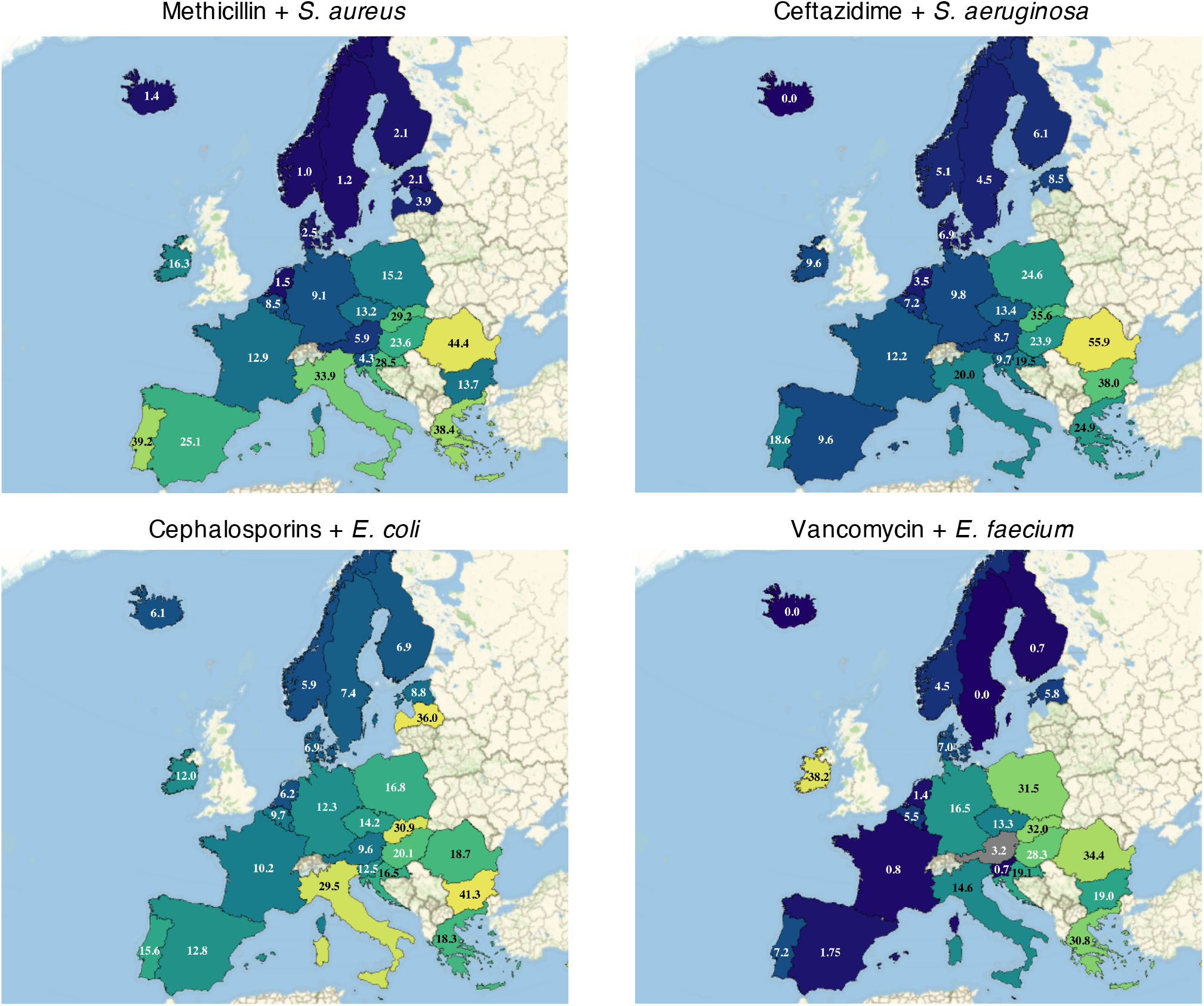
Spatial heterogeneity in resistance levels in Europe for more bug-drug pairs. Percent of isolates resistant to particular antibiotics as reported in the ECDC’s European Antibiotic Resistance Surveillance Network^13,14^ in 2017. Each country is labeled with the resistance level (%). Four example bug-drug pairs are shown: *Staphylococcus aureaus* + methicillin, *Pseudomonas aeruginosa* + ceftazadime, *Eschericia coli* + cephalosporins, *Enterococcus faecium* + vancomycin.

**Figure S3.**
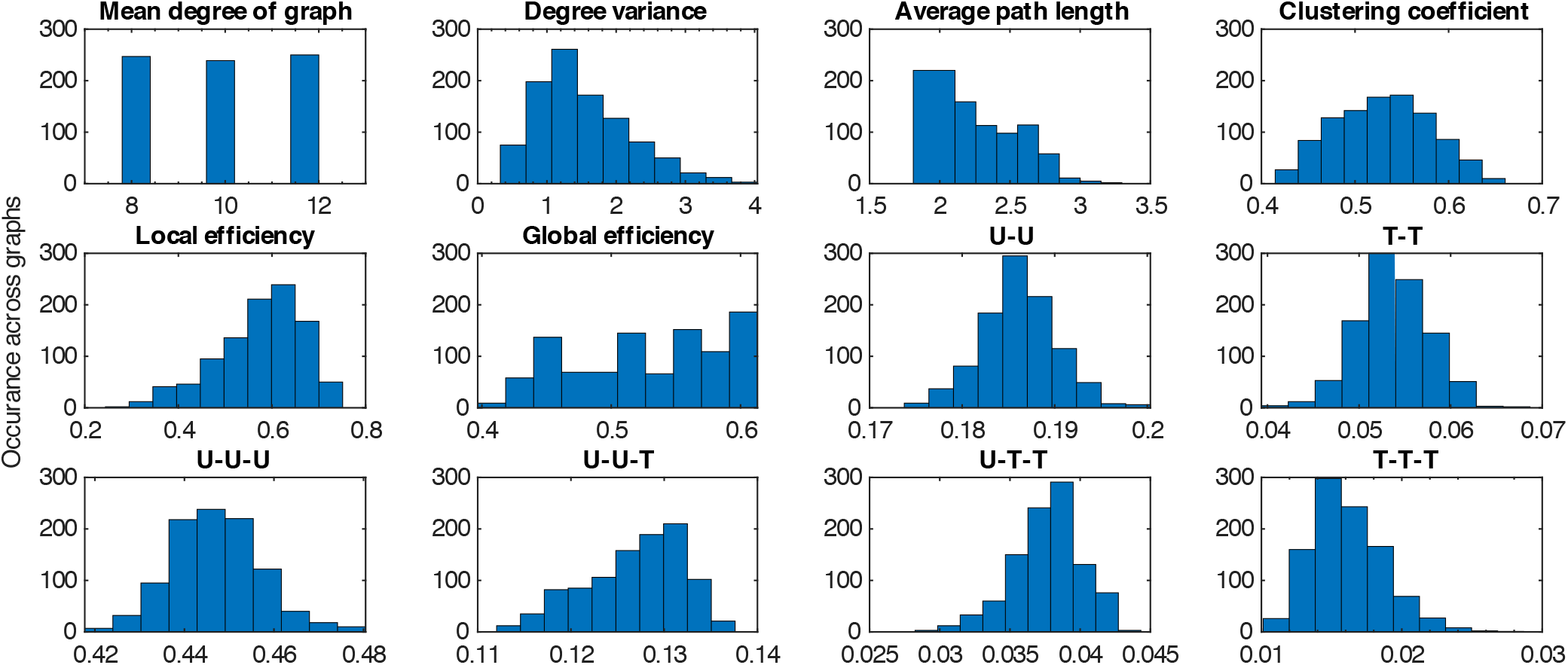
The distribution of graph-theoretic properties of simulated populations used in LASSO regression analysis. Populations were created as random networks of 50 demes using a variant of the Watts-Strogatz algorithm. Treatment was randomly assigned to a portion *ρ* of demes so that an overall desired fraction treated was achieved (here, *ρ* = 0.24). Distributions show properties for 1000 such networks. The first six properties are intrinsic to the network structure, while the latter six describe how treatment is allocated over the network. Quantities of the form X-Y give the proportion of all pairs of connected demes in which one deme has treatment status X and one has treatment status Y (U=untreated, T=treated). Quantities of the form X-Y-Z give the same information for triples of connected demes (order ignored). A description of the methods for creating the networks and for calculating each property is given in the **Methods**.

**Figure S4.**
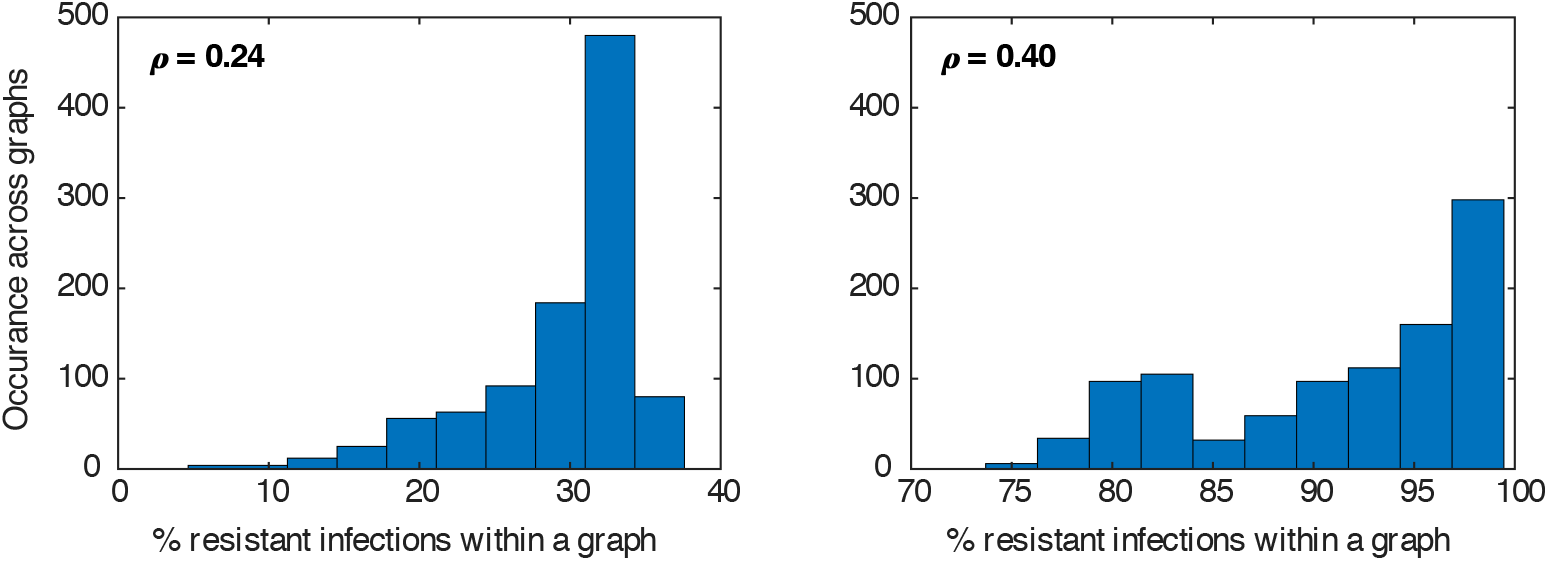
The distribution of the fraction of all infections which are drug-resistant in the simulated populations used in the LASSO regression analysis. Populations were created as random networks of 50 demes using a variant of the Watts-Strogatz algorithm. Treatment was randomly assigned to a portion *ρ* of demes so that an overall desired fraction treated was achieved (on the left *ρ* = 0.24, and on the right *ρ* = 0.4). Infection dynamics were simulated (Eq. (3)) until an equilibrium was reached, at which the proportion of all infections with the drug-resistant strain was recorded. Distributions show results for 1000 such simulations each with a unique random network and treatment allocation. Parameters used were *κ* = 0.25/day, *β* = 0.05/day, *g* = 0.1/day, *ε* = 0.9, *c* = 0.2.

**Figure S5.**
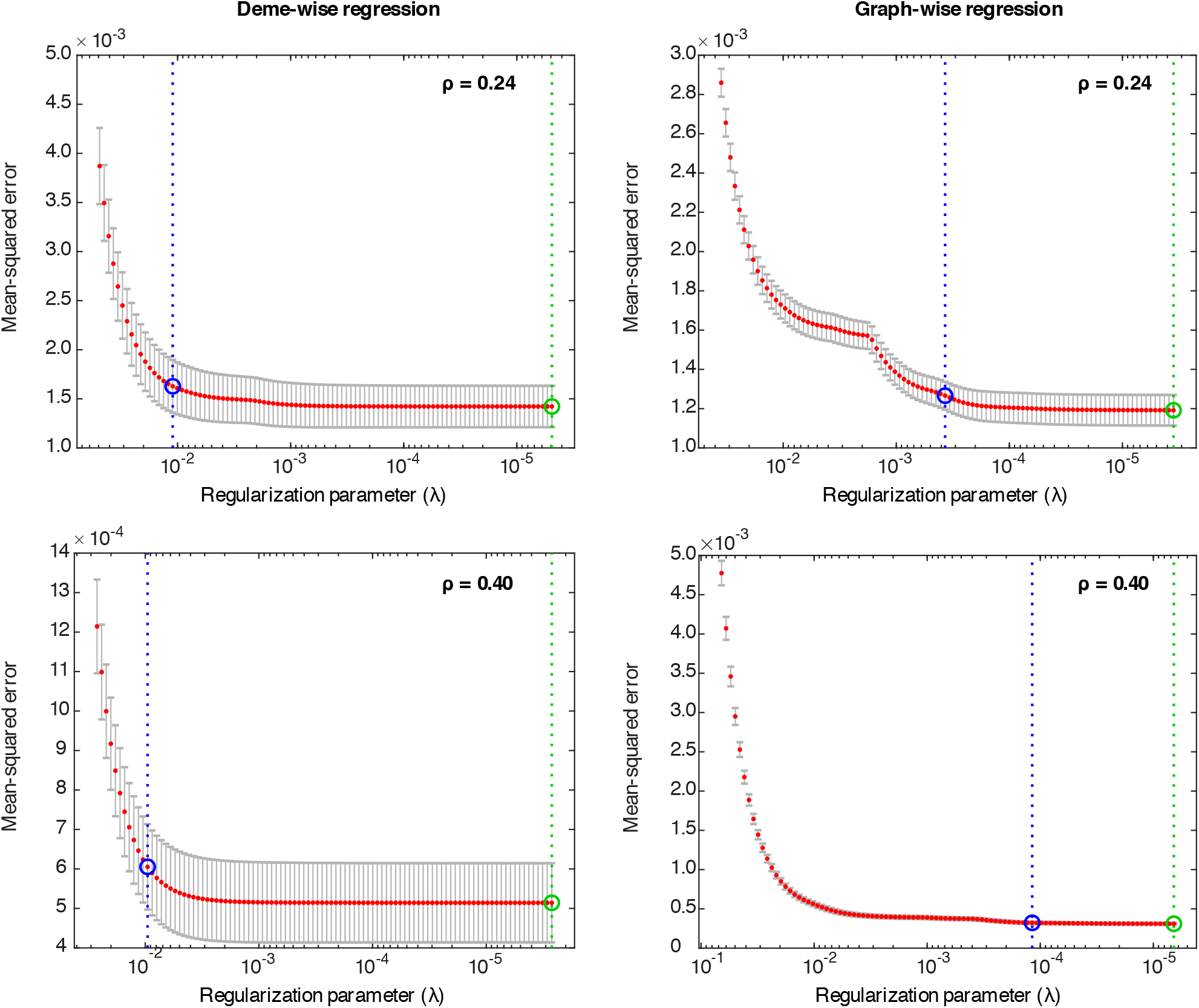
Five-fold cross-validation of LASSO regression models. Mean-squared error of model predictions vs sub-sample of 1/5 of data after model fitting to 4/5 of data, as a function of the internal parameter penalizing the *L*_1_ norm for fitted coefficients (*λ*). Red dots show the mean-squared error of the model, with grey bars showing the standard error of the mean. The green circle/line indicates the value of *λ* that gives the minimum cross-validation error, whereas the blue circle/line locates the *λ* value where the mean-squared error is one standard error above that of the *λ* value with minimum cross-validation error.

**Figure S6.**
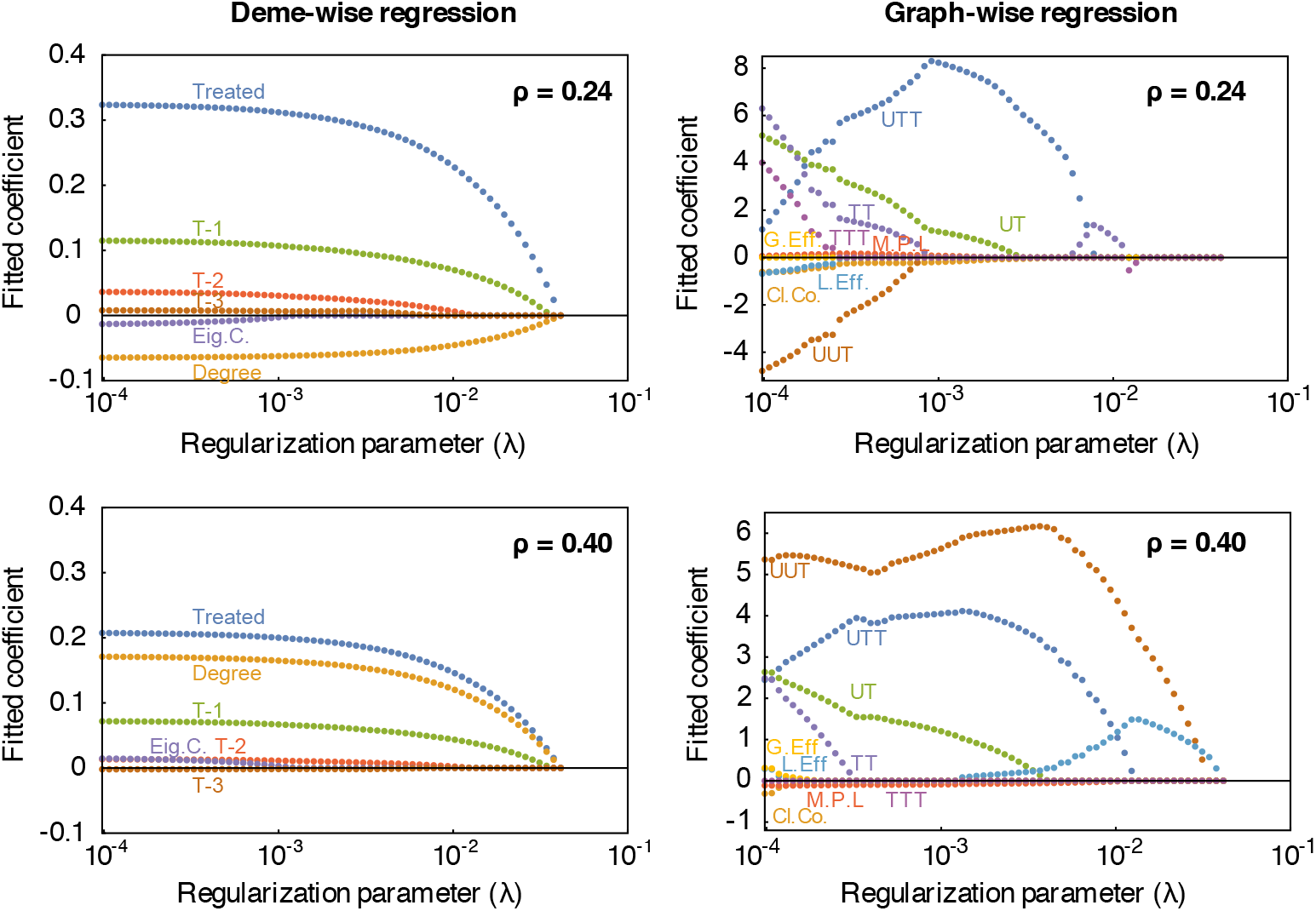
Results of LASSO regression to identify predictors of resistance. Plots show the regression coefficients obtained as a function of the internal constraint on *L_1_* norm of coefficients (“regularization parameter”, *λ*). Each curve is a different predictor variable (described in Table 1). Analysis were separately run to predict deme-level resistance from deme-level properties (left side) or population-level resistance from graph-level properties (right side). Each analysis was conducted for two different levels of drug coverage in the population (either a fraction *ρ* = 0.24 or *ρ* = 0.4 of demes treated). The labels on the curves match the names of the predictors in Table 1.

**Figure S7.**
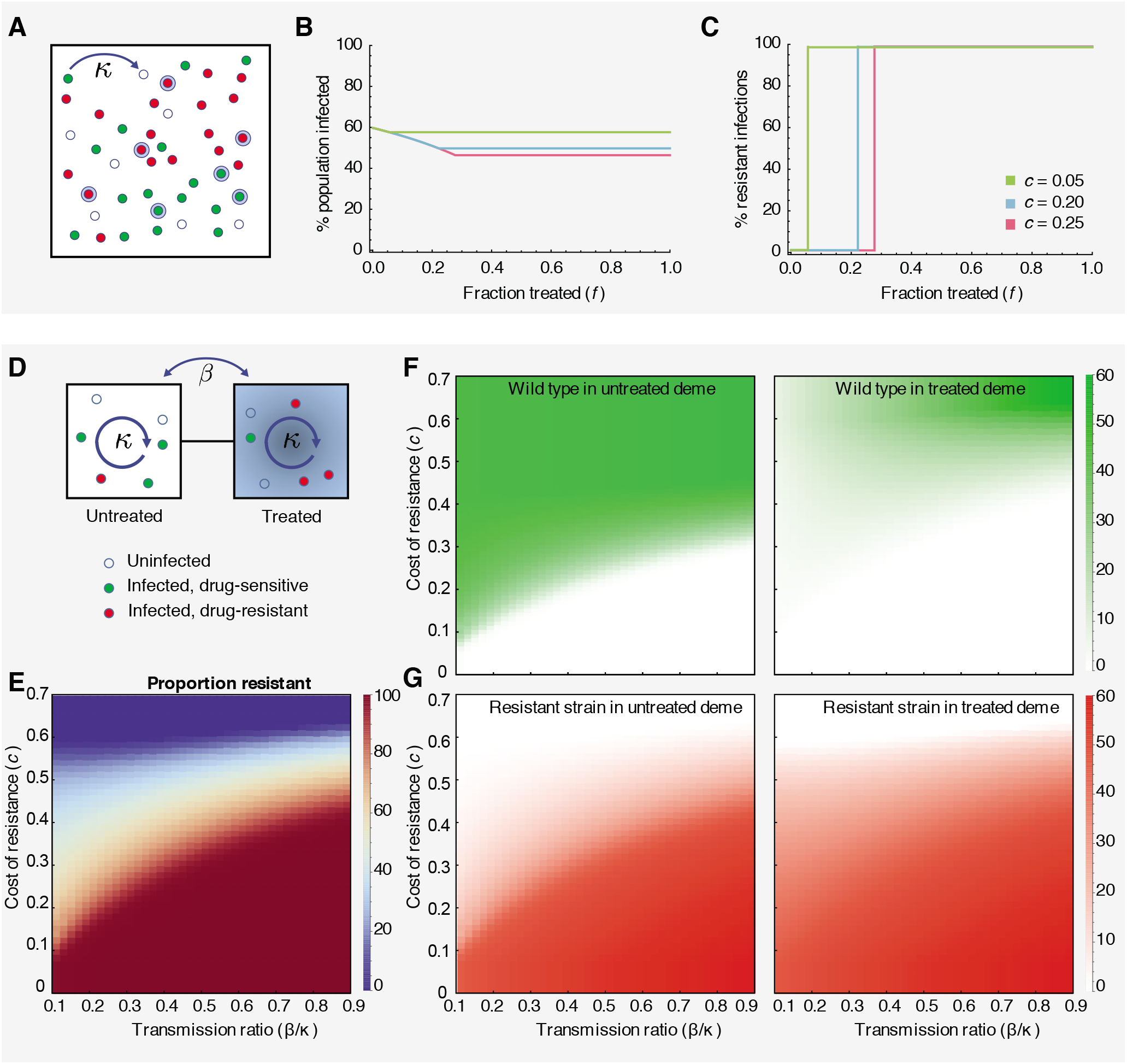
Competition between drug-sensitive and drug-resistant strains in one and two deme populations, when treatment increases recovery rate. **A)** A population of individuals in a single well-mixed deme, in which a fraction *f* will receive drug treatment when infected (blue haloes). Individuals may be uninfected (hollow blue circles), or infected with either the wild-type (green circles) or drug-resistant (red circles) strain. **B)** The total prevalence of infection (wild-type + drug-resistant) as a function of the fraction of treated individuals (*f*) for different costs of resistance (*c*). **C)** The % of infections that are drug-resistant as a function of the fraction of treated individuals (*f*) for different parameters. Infection switches between 0% and 100% resistant when 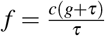. Coexistence never occurs. **D)** Schematic of a two-deme population (left-untreated, right-treated) and the two strains (green-wild type, red-resistant) considered in the model. **E-G)** Each panel shows the infection level (shading) as a function of the relative connectivity between demes (*β / κ*) and the cost of resistance (*c*). **E)** The % of all infections that are drug-resistant strain across the entire population. **F)** The % of individuals in each deme who are infected with the wild type strain. **G)** The % of individuals in each deme who are infected with the resistant strain. For all results, the transmission rate is *κ* = 0.25/day, the recovery rate is *g* = 0.1/day, and the increase in recovery rate due to treatment is treatment efficacy is *τ* = 0.9/day.

**Figure S8.**
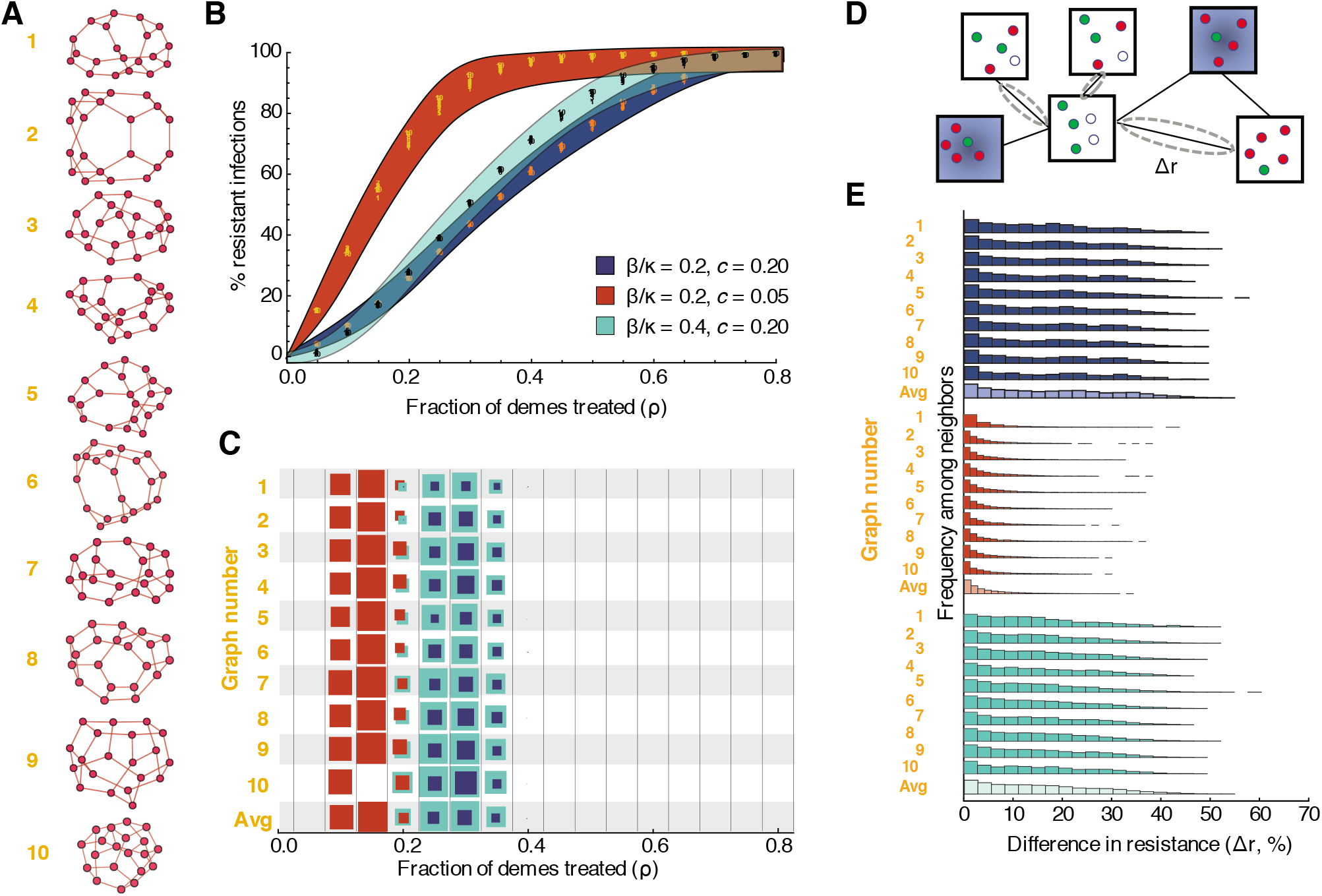
Dynamics of drug-resistant infections in populations consisting of networks of inter-connected demes, when treatment increases recovery rate. **A)** Randomly generated population structures on which infection was simulated. Each node represents a deme (a well-mixed sub-population of individuals), and each edge indicates that infection can spread in either direction between those two demes. Ten example populations were selected out of 1000 total simulated, each with twenty demes randomly connected to three neighbors each, to represent a broad range of outcomes. **B)** Fraction of infections that are resistant in the entire population (*y*-axis) versus fraction of demes treated, *ρ* (*x*-axis). Each color represents a different parameter set (blue background — baseline, red background — lower cost of resistance, teal background — more between-deme connectivity). Numbers show data points for the ten example populations. The colored envelope is created by shading between sigmoidal curves that encompass all the data. **C)** For each population structure shown (*y*-axis) and each treatment level (*x*-axis), the proportion of simulations that resulted in *robust* coexistence between drug-sensitive and drug-resistant strains is shown (by the colored area of the box). Robust coexistence was defined as at least 80% of demes supporting both strains at frequencies above 10%. **D)** Differences in resistance levels (% of all infections that are with the drug resistant-strain) are measured between all pairs of directly-connected untreated demes. **E)** Histograms showing the distribution of pairwise differences in resistance for a given population structure. Lighter shaded histograms combine results from all population graphs. All simulations used kinetic parameters *κ* = 0.25, *g* = 0.1, and *τ* = 0.9, and pooled results from 100 simulations with different random allocation of treatment across demes. Pairwise differences were calculated with 30% treatment.

**Figure S9.**
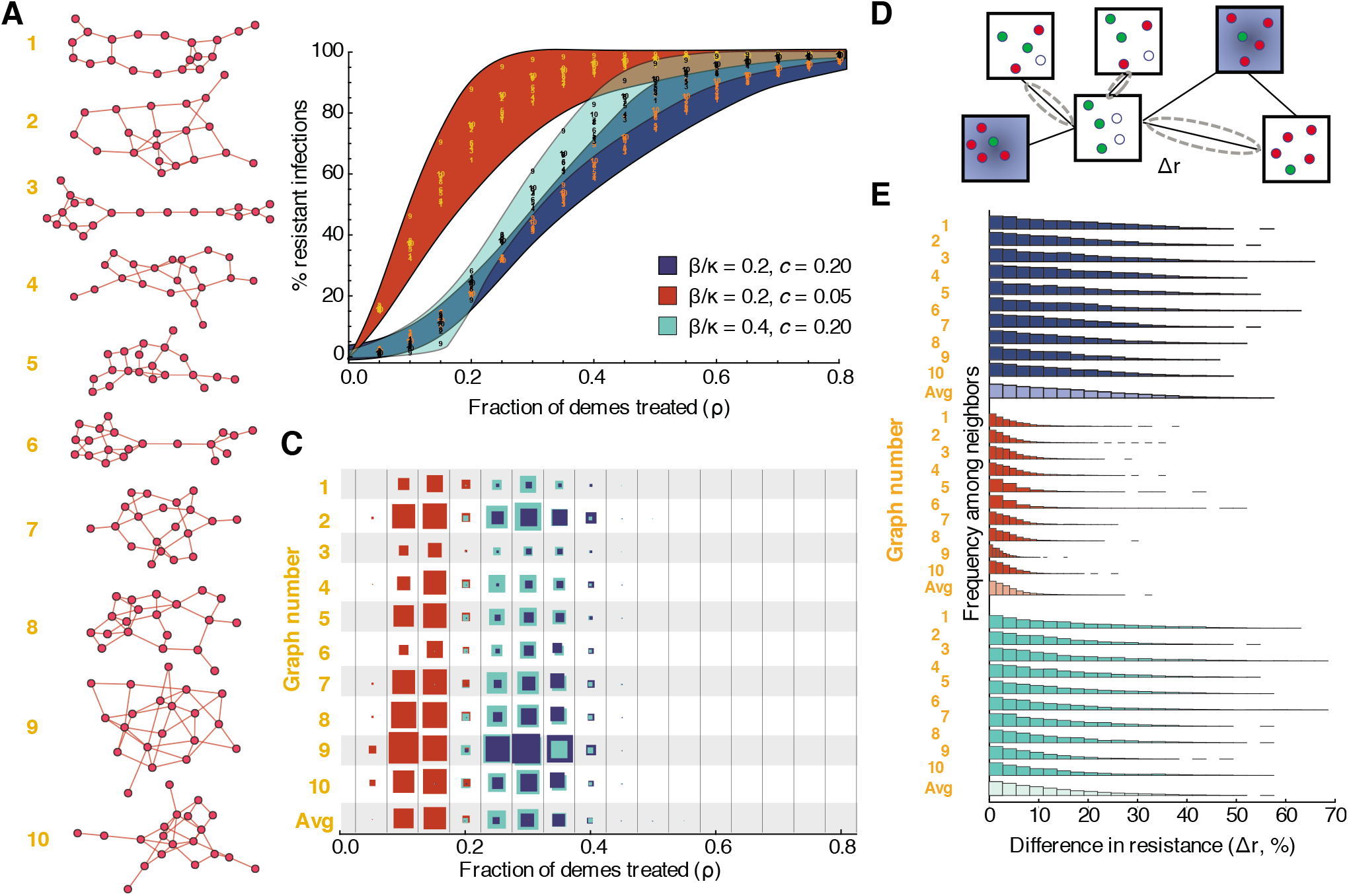
Dynamics of drug-resistant infections in populations consisting of networks of inter-connected demes with heterogeneous connectivity. **A)** Randomly generated population structures on which infection was simulated. Each node represents a deme (a well-mixed sub-population of individuals), and each edge indicates that infection can spread in either direction between those two demes. Each network had twenty demes and the number of neighbors of each deme was drawn from a gamma distribution with mean three and a different variance. Ten example populations were selected out of 1000 total simulated to represent a broad range of variances in connectivity: from graph 1:4.4, 2:5.7, 3:6.2, 4:6.6, 5:6.9, 6:7.2, 7:7.5, 8:8.0, 9:8.6, 10:10.5. **B)** Fraction of infections that are resistant in the entire population (*y*-axis) versus fraction of demes treated, *ρ* (*x*-axis). Each color represents a different parameter set (blue background — baseline, red background — lower cost of resistance, teal background — more between-deme connectivity). Numbers show data points for the ten example populations. The colored envelope is created by shading between sigmoidal curves that encompass all the data. **C)** For each population structure shown (*y*-axis) and each treatment level (*x*-axis), the proportion of simulations that resulted in *robust* coexistence between drug-sensitive and drug-resistant strains is shown (by the colored area of the box). Robust coexistence was defined as at least 80% of demes supporting both strains at frequencies above 10%. **D)** Differences in resistance levels (% of all infections that are with the drug resistant-strain) are measured between all pairs of directly-connected untreated demes. **E)** Histograms showing the distribution of pairwise differences in resistance for a given population structure. Lighter shaded histograms combine results from all population graphs. All simulations used kinetic parameters *κ* = 0.25/day, *g* = 0.1/day, and *ε* = 0.9, and pooled results from 100 simulations with different random allocation of treatment across demes. Pairwise differences were calculated with 30% treatment.

**Figure S10.**
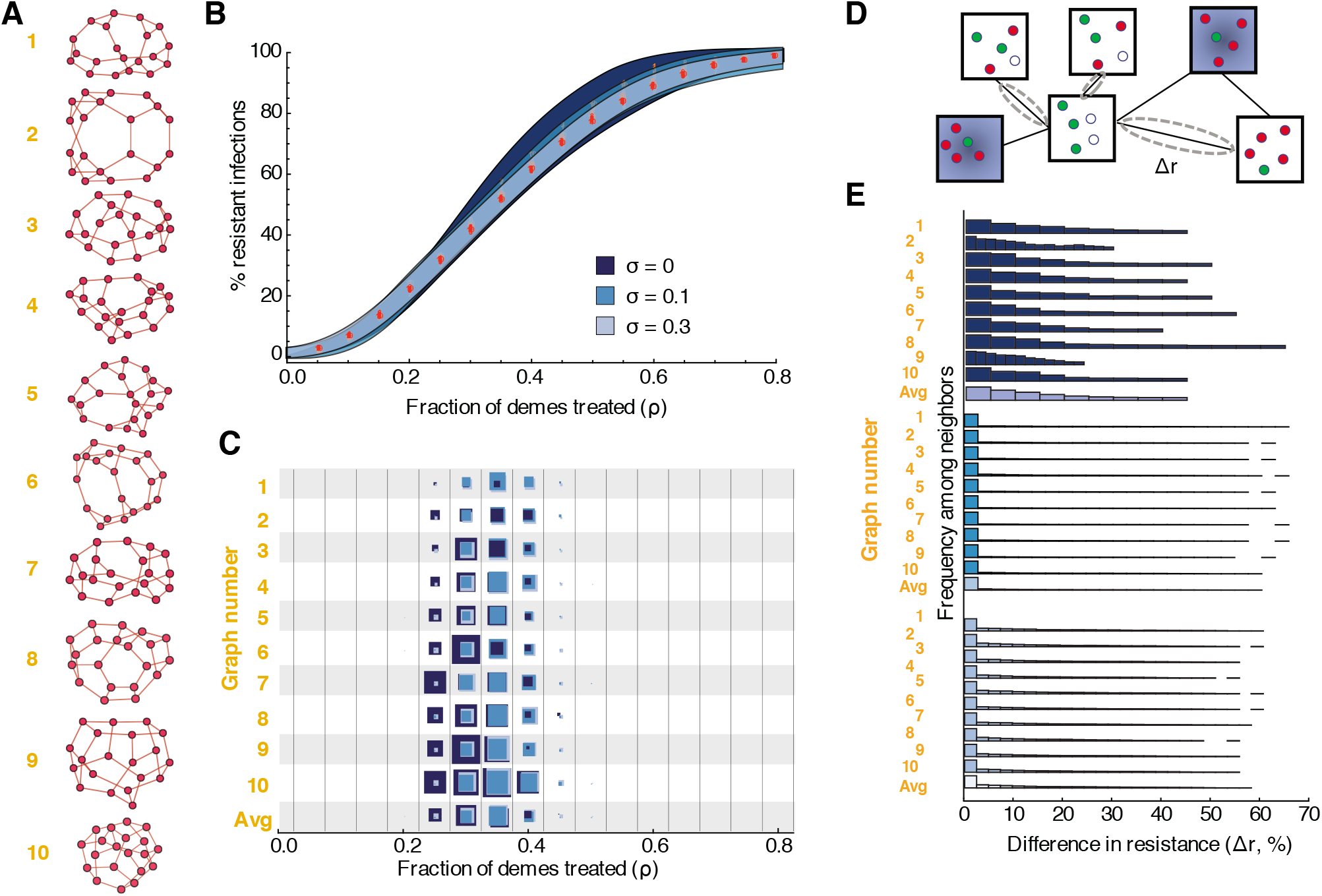
Dynamics of drug-resistant infections in populations consisting of networks of inter-connected demes, with varying deme size. **A)** Randomly generated population structures on which infection was simulated. Each node represents a deme (a well-mixed sub-population of individuals), and each edge indicates that infection can spread in either direction between those two demes. Ten example populations were selected out of 1000 total simulated, each with twenty demes randomly connected to three neighbors each, to represent a broad range of outcomes. Deme sizes were chosen from a normal distribution with coefficient of variation of *σ* =0, 0.1, of 0.3. **B)** Fraction of infections that are resistant in the entire population (*y*-axis) versus fraction of demes treated, *ρ* (*x*-axis). Each color represents a different level of heterogeneity in deme size (n navy blue—uniform sizes (*σ* = 0), medium blue — *σ* = 0.1, light blue — *σ* = 0.3). Numbers show data points for the ten example populations. The colored envelope is created by shading between sigmoidal curves that encompass all the data. **C)** For each population structure shown (*y*-axis) and each treatment level (*x*-axis), the proportion of simulations that resulted in *robust* coexistence between drug-sensitive and drug-resistant strains is shown (by the colored area of the box). Robust coexistence was defined as at least 80% of demes supporting both strains at frequencies above 10%. **D)** Differences in resistance levels (% of all infections that are with the drug resistant-strain) are measured between all pairs of directly-connected untreated demes. **E)** Histograms showing the distribution of pairwise differences in resistance for a given population structure. Lighter shaded histograms combine results from all population graphs. All simulations used kinetic parameters *κ* = 0.25/day, *β* = 0.05/day, *g* = 0.1/day, *c* = 0.2, and *ε* = 0.9, and pooled results from 100 simulations with different random allocation of deme sizes and treatment across demes. Pairwise differences were calculated with 30% treatment.

**Figure S11.**
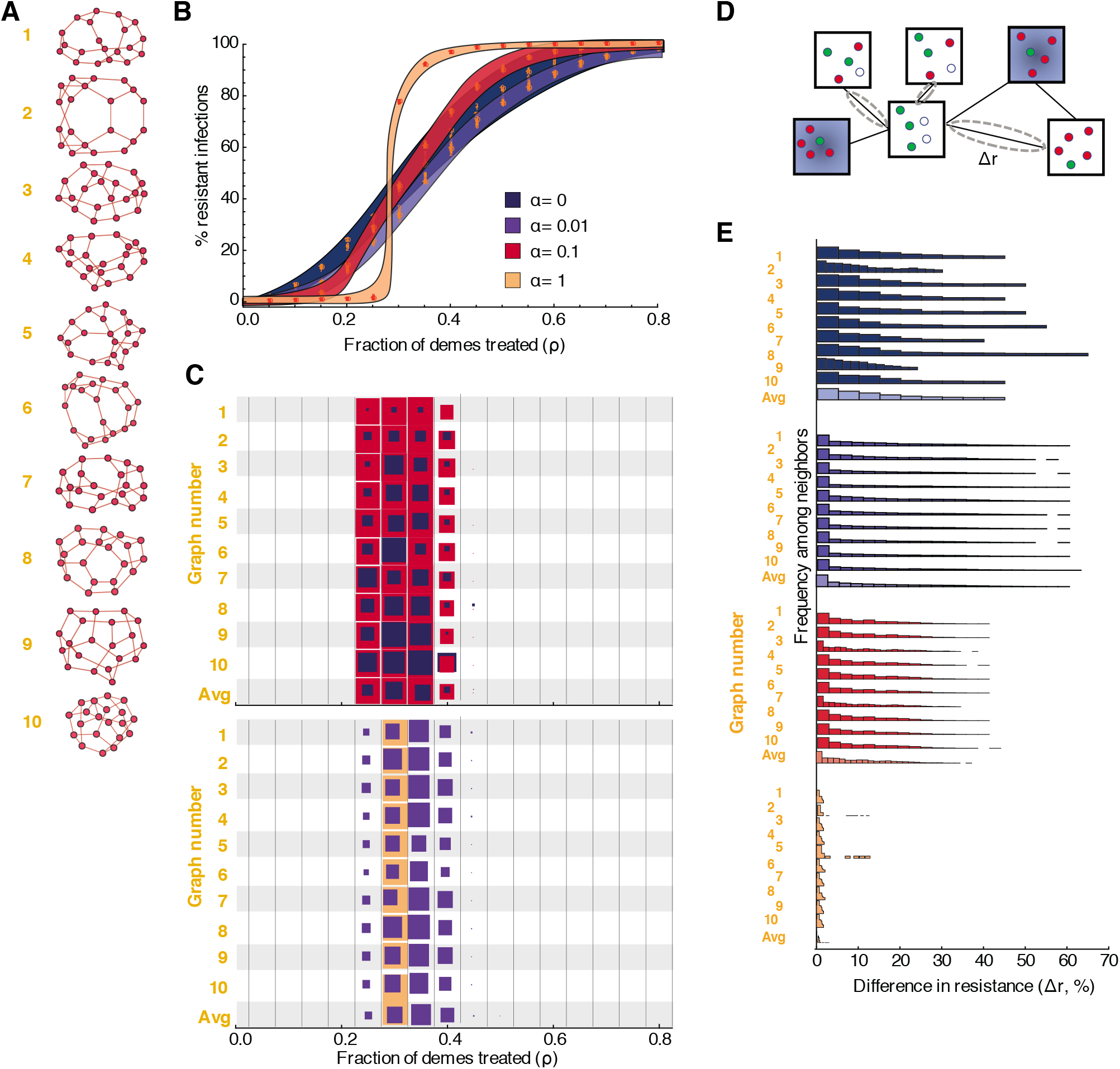
Dynamics of drug-resistant infections in populations consisting of networks of inter-connected demes, with varying levels of background mixing between demes. **A)** Randomly generated population structures on which infection was simulated. Each node represents a deme (a well-mixed sub-population of individuals), and each edge indicates that infection can spread in either direction between those two demes. Each population consisted of twenty demes randomly connected to three neighbors each, and additionally connected with relative weight *α* to all other demes. Ten example populations were selected out of 1000 total simulated to represent a broad range of outcomes. **B)** Fraction of infections that are resistant in the entire population (*y*-axis) versus fraction of demes treated, *ρ* (*x*-axis). Each color represents a different level of background connectivity between all demes (navy blue — no background mixing (*α* = 0), dark blue — *α* = 0.01, medium blue — *α* = 0.1, light blue – *α* = 1). Numbers show data points for the ten example populations. The colored envelope is created by shading between sigmoidal curves that encompass all the data. **C)** For each population structure shown (*y*-axis) and each treatment level (*x*-axis), the proportion of simulations that resulted in *robust* coexistence between drug-sensitive and drug-resistant strains is shown (by the colored area of the box). Robust coexistence was defined as at least 80% of demes supporting both strains at frequencies above 10%. **D)** Differences in resistance levels (% of all infections that are with the drug resistant-strain) are measured between all pairs of directly-connected untreated demes. **E)** Histograms showing the distribution of pairwise differences in resistance for a given population structure. Lighter shaded histograms combine results from all population graphs. All simulations used kinetic parameters *κ* = 0.25/day, *β* = 0.05/day, *g* = 0.1/day, *c* = 0.2, and *ε* = 0.9, and pooled results from 100 simulations with different random allocation of treatment across demes. Pairwise differences were calculated with 30% treatment.

**Figure S12.**
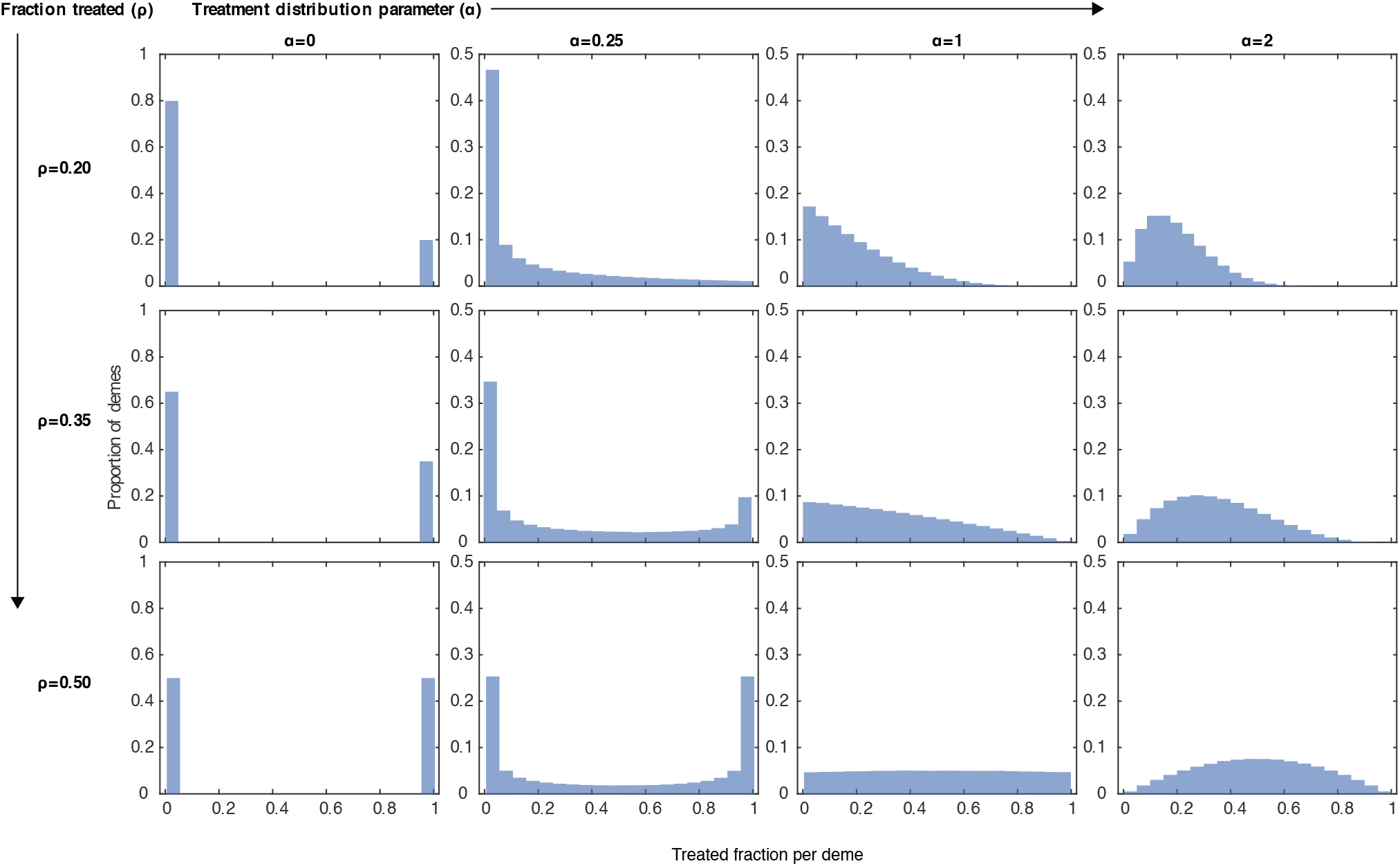
Alternate distributions of treatment across demes. Each panel shows a distribution across demes of the fraction of individuals within the deme receiving treatment if infected. Treatment levels were drawn from Beta distributions with varying mean (*ρ*, rows) and shape parameter (*α*, columns). The left most column corresponds to the results presented in the rest of the paper, where individuals in each deme either have a 0 or 100% chance of being treated if infected.

**Figure S13.**
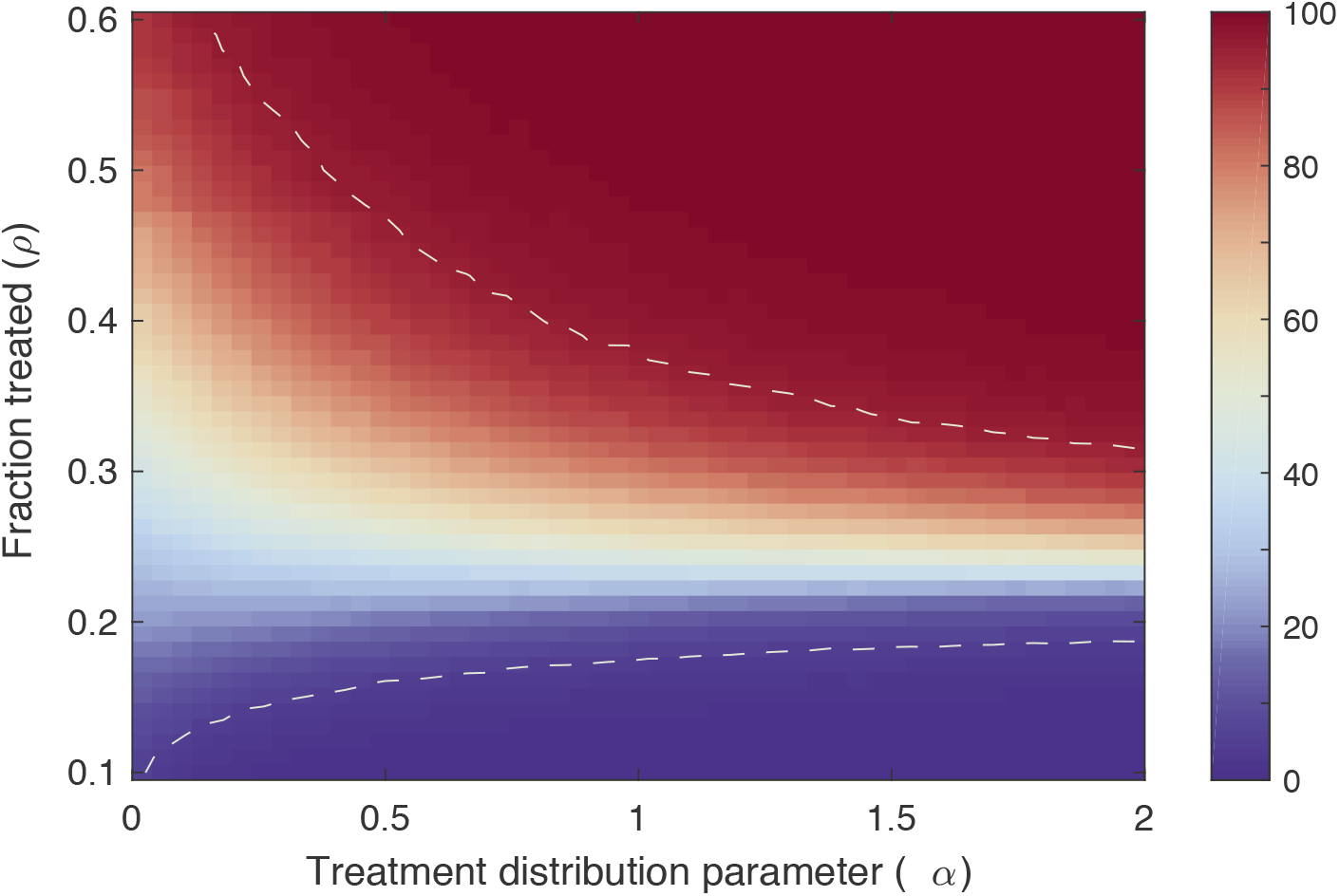
Frequency of drug resistant infections for alternate distributions of treatment across demes. The percent of infections over the whole population that are caused by the the drug resistant strain as a function of the average treatment level across demes (*ρ*, *y*-axis) and the Beta-distribution parameter describing the spread of treatment levels (*α*, *x*-axis). Dotted lines border the region where resistance levels fall between 5% and 95%. Each panel shows a distribution across demes of the fraction of individuals within the deme receiving treatment if infected. Treatment levels were drawn from Beta distributions with varying mean (*ρ*, rows) and shape parameter (*α*, columns). Example treatment distributions are shown in Figure S12. The left most edge of the graph (*α* = 0) corresponds to the results presented in the rest of the paper, where individuals in each deme either have a 0 or 100% chance of being treated if infected. Higher *α* values correspond to more continuous, unimodal treatment distributions with lower variance. The population consisted of twenty demes randomly connected to three neighbors each. Results were averaged over 1000 graphs with with 1000 random treatment allocations for each. For all results, the transmission rates are *κ* = 0.25/day (intra-deme) and *β* = 0.05/day (inter-deme), the recovery rate is *g* = 0.1/day, the cost of resistance is *c* = 0.1, and the treatment efficacy is *ε* = 0.9.

**Figure S14.**
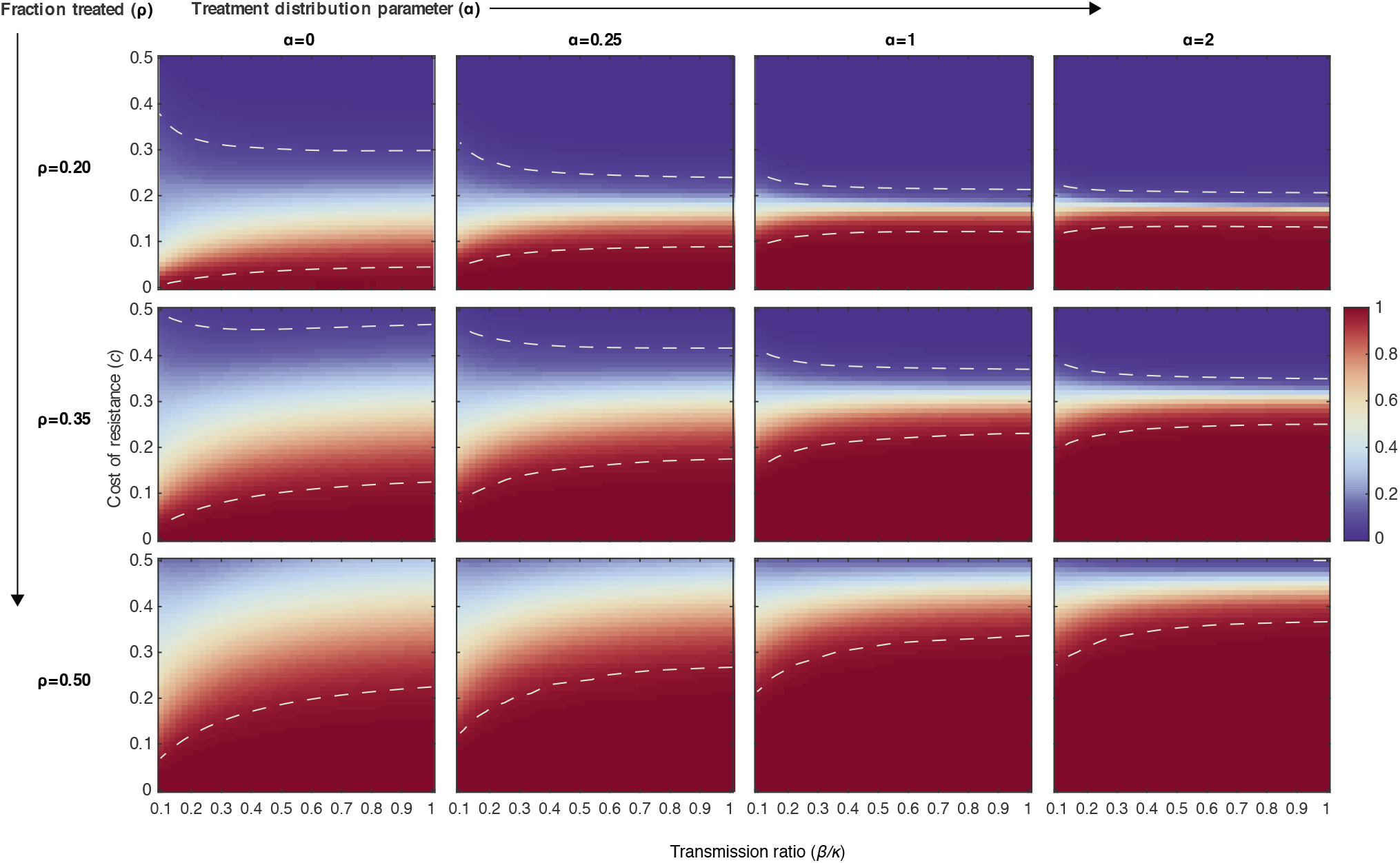
Frequency of drug resistant infections for alternate distributions of treatment across demes. Each panel shows the average fraction of infections that are resistant in the entire population as a function of the relative connectivity between demes (*β / κ*) and the cost of resistance (*c*). Dotted lines border the region where resistance levels fall between 5% and 95%. The mean and variance in the amount of treatment per deme varied between panels and is given by the distributions in Figure S12. The left most column corresponds to the case where individuals in each deme either have a 0 or 100% chance of being treated if infected. Higher *α* values correspond to more continuous, unimodal treatment distributions with lower variance. The population consisted of twenty demes randomly connected to three neighbors each. Results were averaged over 1000 graphs with with 1000 random treatment allocations for each. For all results, the intra-deme transmission rate is *κ* = 0.25/day, the recovery rate is *g* = 0.1/day, and the treatment efficacy is *ε* = 0.9.

**Figure S15.**
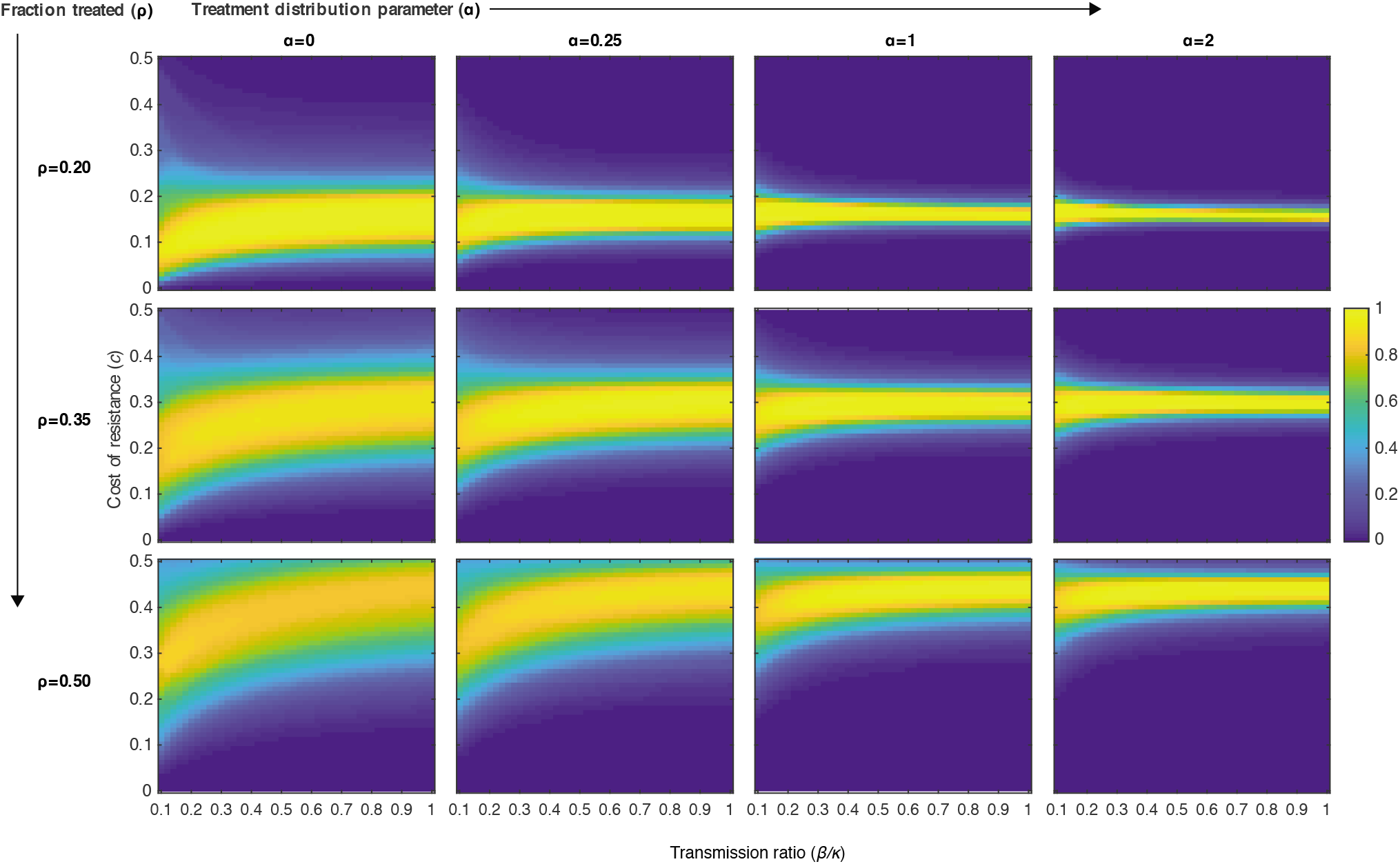
Distributed coexistence for alternate distributions of treatment across demes. Each panel shows the average fraction of demes that supported both strains at frequencies of at least 10%, as a function of the relative connectivity between demes (*β / κ*) and the cost of resistance (*c*). The mean and variance in the amount of treatment per deme varied between panels and is given by the distributions in Figure S12. The left most column corresponds to the case where individuals in each deme either have a 0 or 100% chance of being treated if infected. The population consisted of twenty demes randomly connected to three neighbors each. Results were averaged over 1000 graphs with with 1000 random treatment allocations for each. For all results, the intra-deme transmission rate is *κ* = 0.25/day, the recovery rate is *g* = 0.1/day, and the treatment efficacy is *ε* = 0.9.

**Figure S16.**
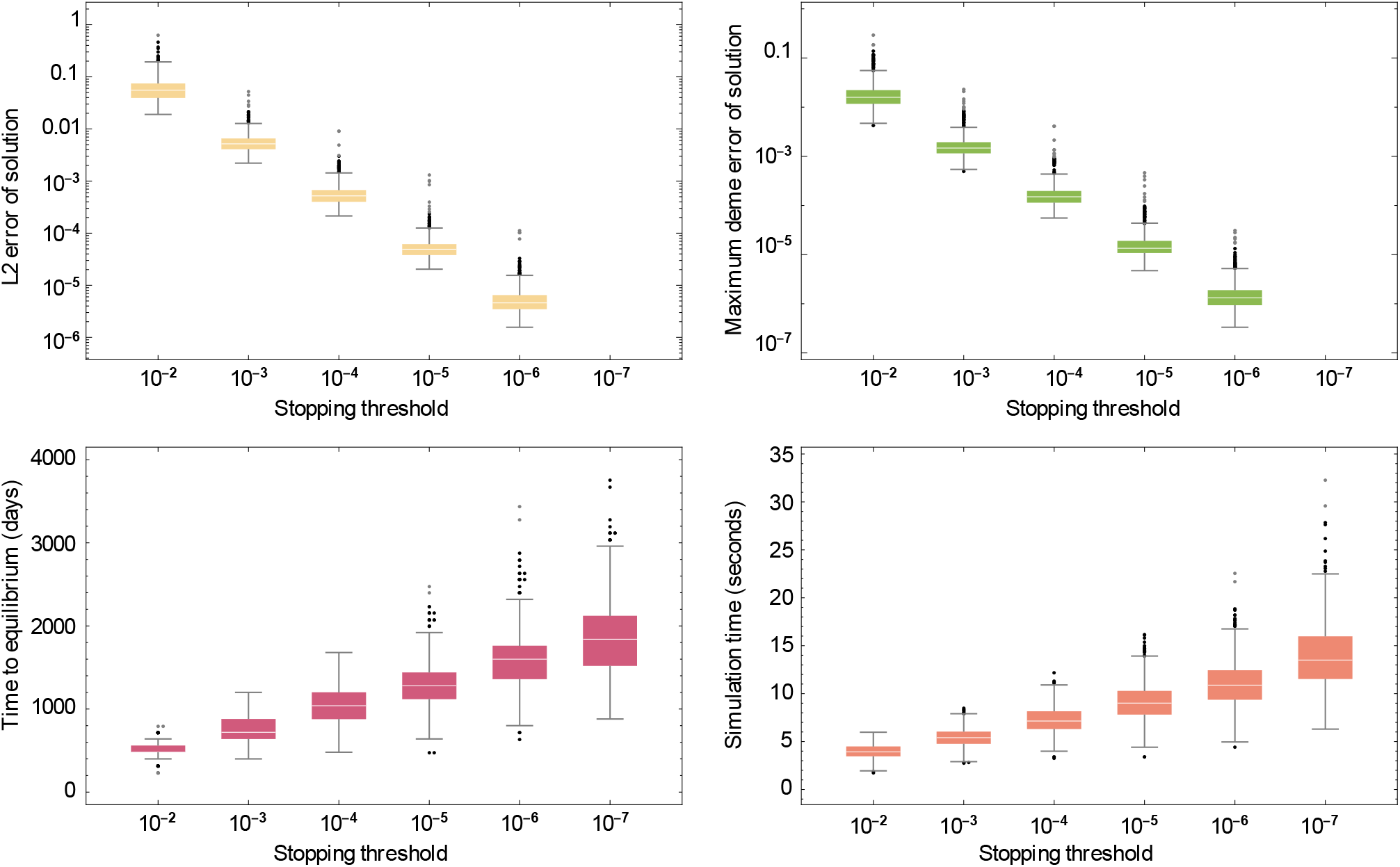
Sensitivity of results to the simulation stopping parameter. We tested the sensitivity of our results to the value of the derivative which we considered “at equilibrium”, defined as the point where the sum of the derivatives of all strains in each deme was less than the stopping parameter divided by the number of demes. **(Top row)** We considered 10^−7^, the smallest value tested, as the “true” solution and measured this solution against other solutions with all parameters held equal, but the stopping parameter increased. Box plots show two representatives of the error from this solution as the stopping parameter is increased up to our parameter 10^−4^ used in the main-text results and beyond: **(left)**, the matrix 2-norm (also known as **L_2_** norm, Frobenius norm, or Euclidean distance). **(Right)**: the maximum demewise error (**L**_∞_ norm). **(Bottom row:** For these same simulations, we show: **(left)**, the time in internal units (simulated days) until equilibrium was reached; **(right)**, the physical computation time of the simulation. The measures of error for our chosen parameter are small given the advancement in computation time afforded by that choice.

**Figure S17.**
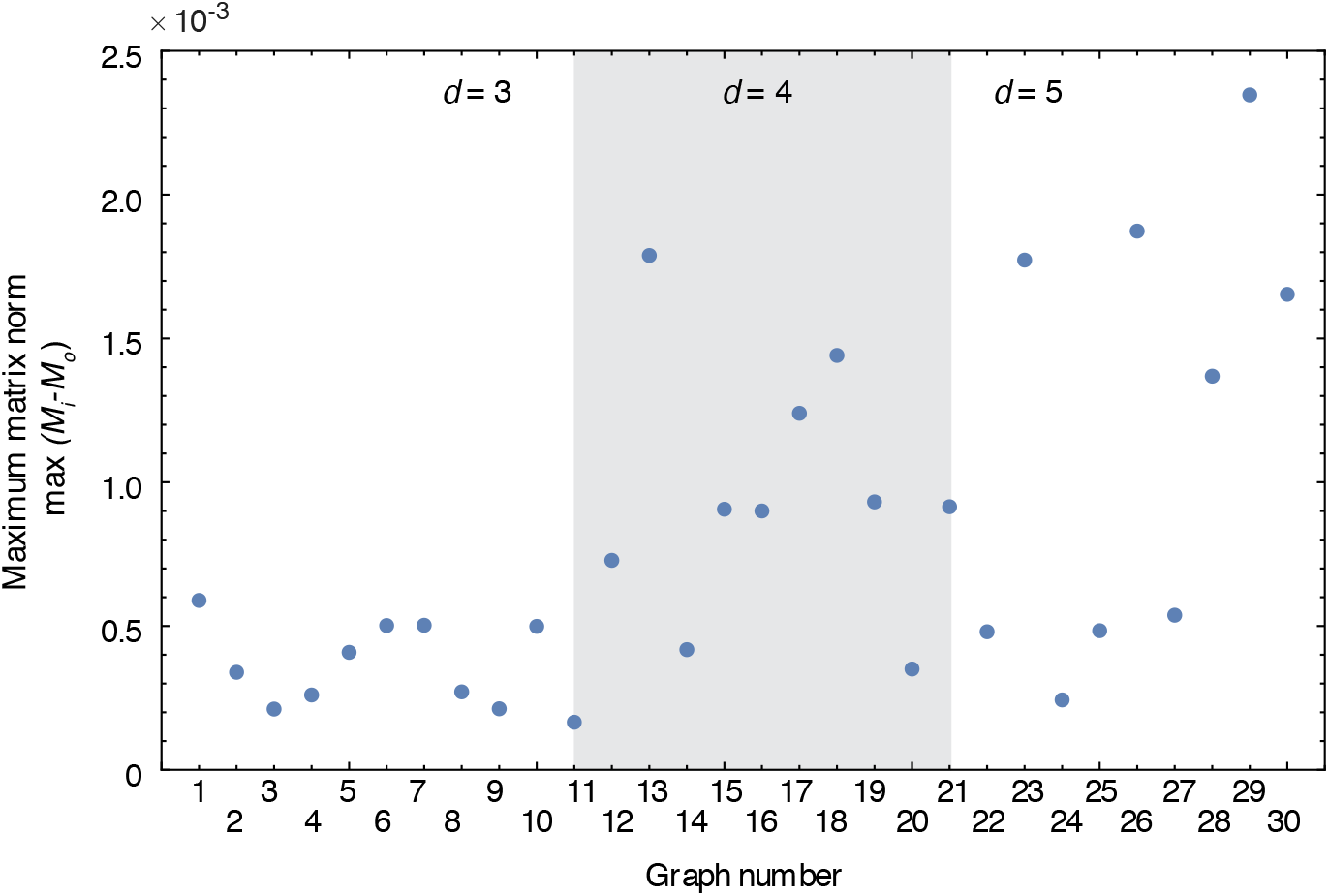
Difference between equilibrium values obtained for infection dynamics in multi-deme systems as initial conditions vary. For 10 different random regular graphs with 20 demes and degree (*d*) 3, 4, or 5, treatment was allocated randomly across demes with an overall proportion treated *ρ* = 0.35. For each graph-treatment allocation combination, 100 different sets of initial conditions (levels of resistant and sensitive infections in each deme) were chosen uniformly at randomly, and infection dynamics were run until an equilibrium was reached as described in the Methods. The maximum difference (in matrix 2-norm, also known as **L_2_** norm, Frobenius norm, or Euclidean distance) was calculated between the equilibria seen in any of 100 trials and the equilibrium values used for results reported in the main text. Very low maximum norm values suggest there is a single stable equilibrium value for each parameter set.

## Notes

### Competing Interest Statement

The authors have declared no competing interest.

